# Adintoviruses: An Animal-Tropic Family of Midsize Eukaryotic Linear dsDNA (MELD) Viruses

**DOI:** 10.1101/697771

**Authors:** Gabriel J. Starrett, Michael J. Tisza, Nicole L. Welch, Anna K. Belford, Alberto Peretti, Diana V. Pastrana, Christopher B. Buck

## Abstract

Polintons (also known as Mavericks) were initially identified as a widespread class of eukaryotic transposons named for their hallmark type B DNA *pol*ymerase and retrovirus-like *int*egrase genes. It has since been recognized that many polintons encode possible capsid proteins and viral genome-packaging ATPases similar to those of a diverse range of double-stranded DNA (dsDNA) viruses. This supports the inference that at least some polintons are viruses that remain capable of cell-to-cell spread. At present, there are no polinton-associated capsid protein genes annotated in public sequence databases. To rectify this deficiency, we used a data-mining approach to investigate the distribution and gene content of polinton-like elements and related DNA viruses in animal genomic and metagenomic sequence datasets. The results define a discrete family-like clade of animal-specific viruses with two genus-level divisions. We suggest the family name *Adintoviridae,* connoting similarities to *ad*enovirus virion proteins and the presence of a retrovirus-like *int*egrase gene. Although adintovirus-class PolB sequences were detected in datasets for fungi and various unicellular eukaryotes, sequences resembling adintovirus virion proteins and accessory genes appear to be restricted to animals. Degraded adintovirus sequences are endogenized into the germlines of a wide range of animals, including humans.

## Introduction

Analyses based on conserved protein structural features have increasingly revealed commonalities between families of eukaryotic viruses with double-stranded DNA (dsDNA) genomes. A current model places a loosely defined group known as polinton-like viruses at the center of a network of evolutionary relationships (Koonin, Dolja et al. 2015, Koonin, Krupovic et al. 2015). Polintons (also known as Mavericks) are defined by the presence of a type B DNA *pol*ymerase (PolB) and a retrovirus-like *int*egrase gene. Although polintons were first recognized as transposons, the observation that many of them encode predicted virion proteins supports the proposal that most elements initially designated as polinton transposons are actually integrated proviruses that may remain capable of infectious cell-to-cell spread (Krupovic, Bamford et al. 2014, Krupovic and Koonin 2015).

Adenoviruses, poxviruses, and baculoviruses are familiar groups of animal-tropic viruses that encode genes distantly similar to polinton PolB and virion proteins (Koonin, Dolja et al. 2015). An emerging group of viruses known as virophages, which are named for their ability to parasitize megaviruses that infect unicellular eukaryotes, also encode polinton-like PolB and virion protein genes as well as, in some cases, retrovirus-like integrase genes. (Duponchel and Fischer 2019).

Although polintons have been widely recognized in animal genomics and transcriptomics datasets (Krupovic, Bamford et al. 2014), the proposed capsid genes of these elements are not currently annotated in public sequence databases. This has led to confusion. For instance, a recent study detected two “Maverick transposons” in insect cell cultures but failed to annotate the capsid genes that identify them as likely viruses (Geisler 2018). In another example, a set of classic polinton PolB gene fragments detected in mouse fecal samples appear in GenBank with annotations incorrectly indicating that they are parvovirus structural proteins (Williams, Che et al. 2018). A primary goal of this study is to develop a coherent classification system for animal-tropic viruses with polinton-like genes and to facilitate further discovery by rendering annotated examples of these viruses searchable in public databases.

## Results

### Classification of animal-associated contigs with polinton-like PolB genes

TBLASTN searches using the inferred virion maturational protease (Adenain) of an arbitrarily chosen *Parasteatoda* spider contig (AOMJ02256338) identified hundreds of >10kb contigs of interest in NCBI’s whole genome shotgun (WGS) and transcriptome shotgun assembly (TSA) databases, as well as in *de novo* assemblies of various datasets of interest from the Sequence Read Archive (SRA). In animal datasets, a great majority of the larger adenain-bearing elements were found to encode either an archetypal polinton-like PolB (pfam03175) or a divergent PolB <30% identical to the pfam03175 type. Both PolB types encode a distinctive N-terminal domain with predicted structural similarity to the ovarian tumor superfamily of ubiquitin-specific proteases (OTU). Adenovirus PolB sequences lack the OTU domain. In this study, we refer to the OTU-pfam03175 PolB class as Alpha and the second OTU-PolB class as Beta. In RepBase https://www.girinst.org/, polinton groups 1, 2, 3, 4, and 9 each contain both Alpha and Beta PolB genes. Alpha PolB genes have previously been binned with hybrid virophages, ungrouped polinton-like viruses, and Polintons group 2, while Beta PolB genes have been binned with ungrouped polinton-like viruses, plant and fungal mitochondrial plasmids, and Polintons group 1 (Moriyama, Terasawa et al. 2008, Yutin, Raoult et al. 2013, Yutin, Kapitonov et al. 2015, Yutin, Shevchenko et al. 2015).

In BLASTP searches, Alpha PolB sequences give strong hits (E-values ~1e-60) for an emerging group of bipartite parvovirus-like viruses called bidnaviruses or bidensoviruses (Krupovic and Koonin 2014). Use of the DELTA-BLAST algorithm (Boratyn, Schaffer et al. 2012) yields stronger hits (E-values <1e-100) for adenoviruses. Beta PolB sequences typically do not yield bidnavirus hits in BLASTP searches and instead give moderate hits (E-value ~1e-15) for the PolB proteins of megaviruses (e.g., Faustovirus and Klosneuvirus) as well as various bacteriophages (Figure 1). Neither of the two PolB classes detects known virophage PolB sequences in BLASTP or DELTA-BLAST searches.

**Figure 1:**
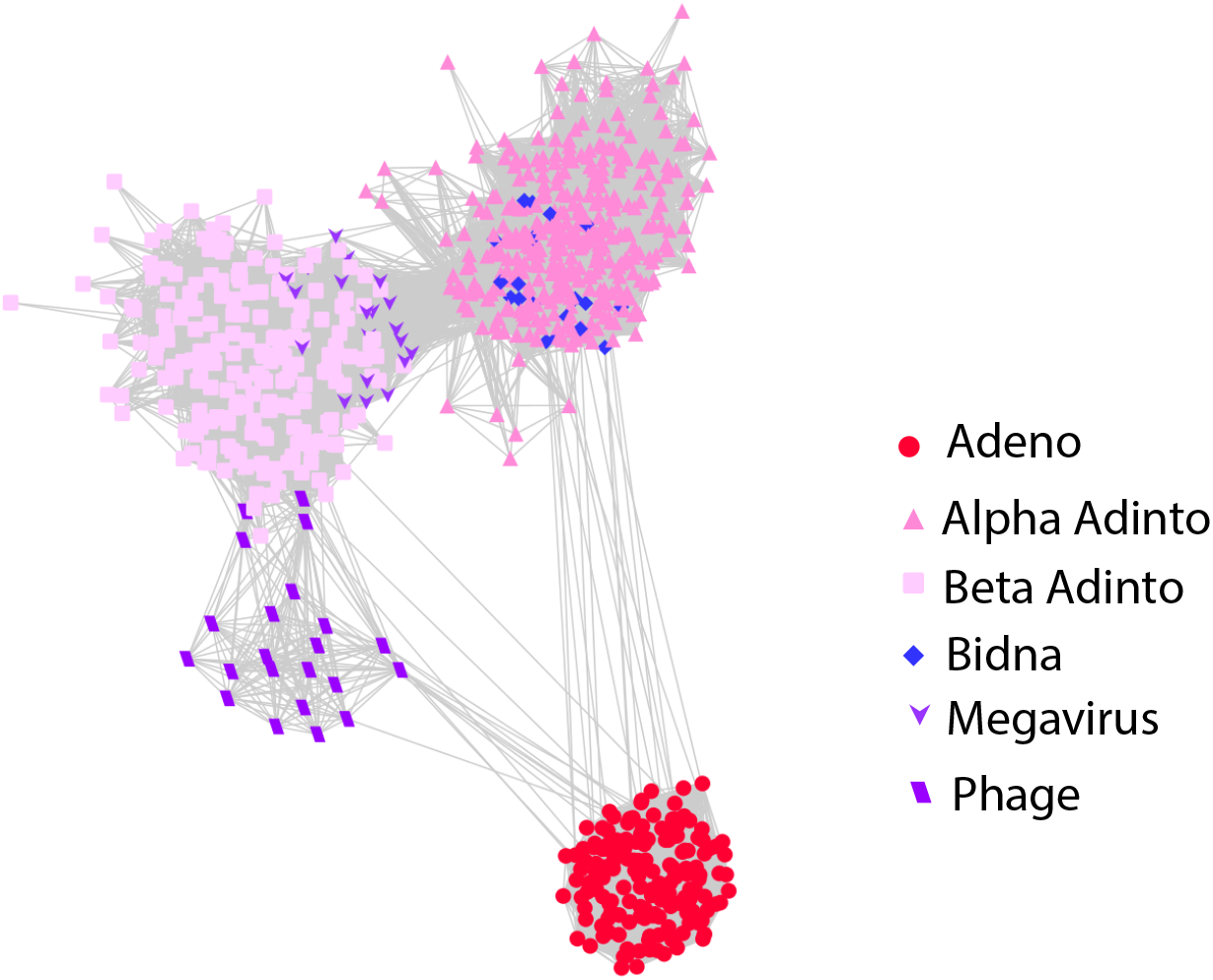
PolB BLASTP relationships. PolB protein sequences were subjected to all-against-all sequence similarity network analysis with a BLASTP E-value cutoff of 1e-06. Figure supplement 1: an interactive version of Figure 1 that can be viewed using Cytoscape software https://cytoscape.org Figure supplement 2: sequence compilations for PolB and other proteins (fasta format, zip compressed) Figure supplement 3: network analysis of Hexon and Penton proteins

In addition to Adenain and PolB, nearly all >10 kb contigs from the WGS and TSA surveys encode a retrovirus-like integrase (protein family rve) as well as a protein similar to a group of FtsK/HerA-type nucleoside triphosphatases (FtsK) that are thought to mediate the packaging of viral genomes into virions (Iyer, Makarova et al. 2004).

Alignments of selected contigs back to parent read datasets showed that coverage depth fell to zero near the ends of some contigs. An example is shown graphically in Figure 2 Figure supplement 2. In some cases, such as a *Mayetiola destructor* (barley midge) read dataset, a single predominant apparently free-ended sequence could be assembled but the dataset also contained a range of lower-coverage variant reads near the termini, some of which extended into inverted terminal repeats (ITRs) and host genomic DNA sequences. The observation suggests that the integrase gene is functional and mediates integration events akin to those observed in virophages that encode rve integrases (Fischer and Hackl 2016).

**Figure 2:**
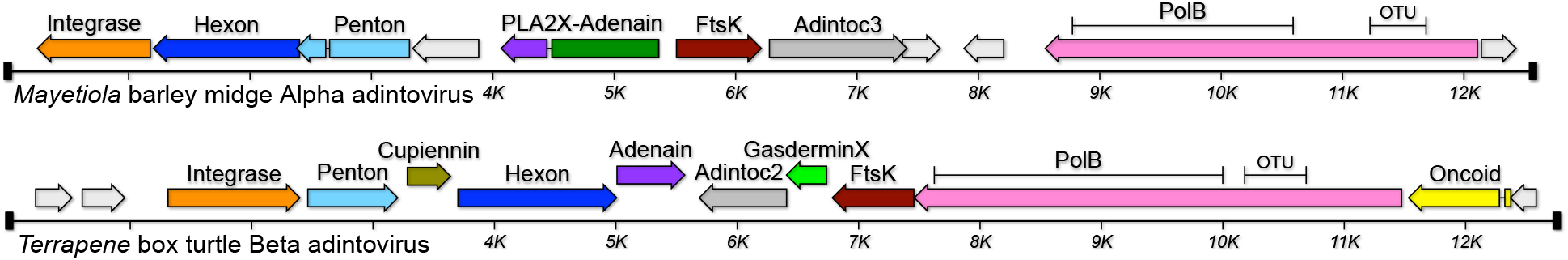
Genome maps of two representative adintoviruses. Figure supplement 1: accession numbers and full Linnaean designations of animal hosts (MS Excel table). Figure supplement 2: graphical examples of the gene-annotation process. Figure supplement 3: graphical maps of additional adintoviruses. Figure supplement 4: annotated nucleotide maps of adintoviruses and related viruses (GenBank-formatted text file).

Based on the similarities to *ad*enoviruses and the presence of *int*egrase and *virus* genome-packaging genes we suggest that this group of animal-associated elements could be referred to as “adintoviruses.” Maps of reference adintoviruses are shown in Figure 2.

HHpred searches confirmed the presence of ORFs with high-probability predicted structural similarity to the double-jellyroll major capsid proteins (Hexons) and single-jellyroll vertex minor capsid proteins (Pentons) of adenoviruses, virophages, megaviruses, or poxviruses (see Figure 2 Figure supplement 2 for illustrated examples of annotation methods). As expected, contigs with Beta PolB genes encode Hexon and Penton proteins that occupy discrete clusters that encompass the *Terrapene* Beta adintovirus cognates (Figure 1 Figure supplement 3). Although most contigs with Alpha PolB genes encode Hexon and Penton proteins that cluster with the *Mayetiola* cognates, some Alpha PolB contigs unexpectedly encode virion proteins that are interspersed within the *Terrapene* cluster. Similar results were observed in analyses using traditional phylogenetic trees. The results suggest the existence of distinct Alpha and Beta adintovirus lineages, but with some examples reflecting horizontal transfer of virion protein operons from the Beta PolB lineage into the Alpha PolB lineage. We have previously proposed a similar intra-family horizontal gene transfer scenario for some species of polyomaviruses (Buck, Van Doorslaer et al. 2016).

### Other adintovirus genes

Adintoviruses encode three classes of proteins with predicted structures resembling known membrane-active proteins. A previously noted class (Yutin, Raoult et al. 2013) is similar to the phospholipase A2 (PLA2) domain of parvovirus VP1 virion proteins (Figure 2 Figure supplement 2). In parvoviruses, the domain is thought to be involved in membrane disruption during the infectious entry process. The PLA2-like genes, which are characteristic of *Mayetiola*-class (Alpha) virion protein operons, include a C-terminal domain similar to adenovirus virion core protein ten (pX). We suggest the gene name PLA2X.

Beta adintoviruses, as well as Alpha PolB adintoviruses with *Terrapene*-class (Beta) virion protein operons, encode homologs of the C-terminal regulatory domain of gasdermins, a group of pore-forming proteins that serve as executioners in pyroptosis (a form of inflammatory programmed cell death)(Dubois, Sorgeloos et al. 2019). Like PLA2X, adintovirus gasdermin homologs typically encode a pX-like domain near the C-terminus. Apparent homologs of a membrane-active spider venom protein known as cupiennin were also observed in Beta-class virion protein operons. The pairing of hallmark Beta-class virion accessory genes (GasderminX, Cupiennin) with a subset of Alpha PolB adintoviruses (Figure 2 Figure supplement 3) supports the hypothesis that some adintovirus species arose through chimerization between the Alpha and Beta adintovirus lineages.

Some classes of predicted protein sequences were conserved among adintoviruses but did not show clear hits for known proteins in BLASTP or HHpred searches. We assigned these groups of adintovirus-conserved proteins of unknown function numbered “Adintoc” names.

Small DNA tumor viruses (adenoviruses, polyomaviruses, and papillomaviruses (Pipas 2019)) encode proteins harboring conserved LXCXE motifs that that are known to engage cellular retinoblastoma (Rb) and related tumor suppressor proteins (de Souza, Iyer et al. 2010). Adenovirus E1A, papillomavirus E7, polyomavirus LT, and parvovirus NS3 oncoproteins typically encode the Rb-binding motif just upstream of a consensus casein kinase 2 acceptor motif ((ST)XX(DE)). Some oncogenes, such as E1A, encode an additional conserved region ((DEN)(LIMV)XX(LM)(FY)), referred to as CR1, that binds the groove containing the A and B cyclin folds within the Rb pocket domain (Pipas 1992, Gouw, Michael et al. 2018). In general, these predicted Rb-interacting motifs are adjacent to potential zinc- or iron-sulfur-binding motifs (typically, paired CXXC). Open reading frames encoding combinations of these short linear motifs were observed in adintovirus contigs. We refer to these predicted proteins, which typically occupy a region upstream of the PolB gene, as “Oncoid” genes, conjnoting their similarities to the known oncogenes of small DNA tumor viruses. Adintovirus homologs of anti-apoptotic proteins, such as Bcl2 and IAP, were also observed (Figure 2 Figure supplement 3).

### Distribution of adintovirus-like PolB sequences in eukaryotic WGS datasets

The conserved catalytic core PolB sequences of either the *Mayetiola* barley midge Alpha adintovirus or *Terrapene* box turtle Beta adintovirus were used separately as baits in TBLASTN searches of WGS databases for eukaryotes. Retrieved protein sequences were trimmed to 80% similarity and subjected to clustering with an alignment score threshold of 60 (Shannon, Markiel et al. 2003, Li and Godzik 2006, Huang, Niu et al. 2010, Fu, Niu et al. 2012, Zallot, Oberg et al. 2018). The clustering segregated away Beta adintovirus-like PolB sequences encoded by plant and fungal mitochondria (e.g., EU365401, AF061244). The filtered sequences were subjected to phylogenetic analyses (Figure 3).

**Figure 3:**
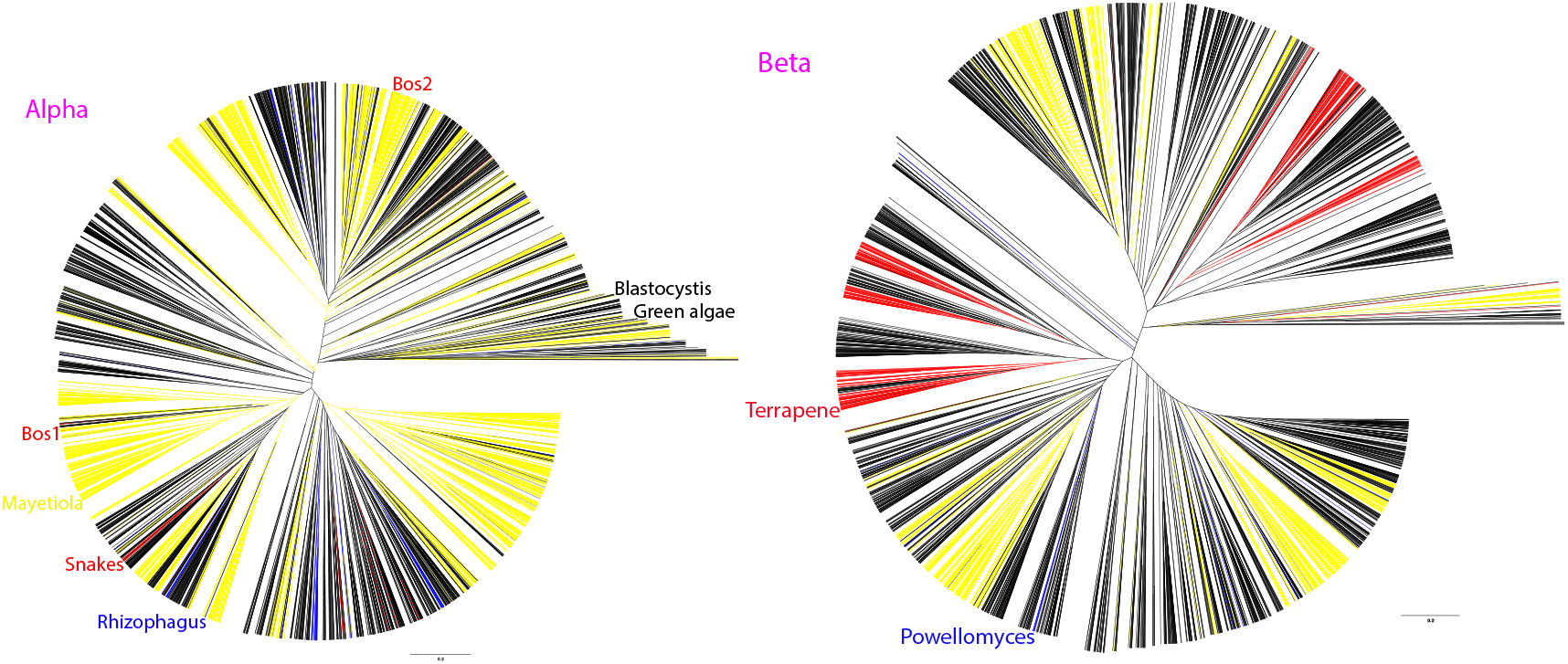
Phylogenetic trees comprised of WGS hits for Alpha or Beta adintovirus PolB sequences (left and right panels, respectively). Hits from insect datasets are colored yellow, hits from tetrapod datasets are red and fungus-associated hits are blue. All other types of eukaryotes are represented by black lines. Annotated branches show two Alpha adintovirus sequences associated with bovine (*Bos*) lung samples clustering with adintovirus sequences from insect datasets, suggesting an environmental insect source. In contrast, exemplar *Mayetiola* and *Terrapene* adintoviruses cluster with sequences found in other insect or terrestrial vertebrate datasets, respectively. Similarly, adintovirus PolB-like sequences from *Powellomyces* and *Rhizophagus* fungi cluster with sequences from other types of fungi. Figure supplements 1 and 2: interactive Nexus-format tree files that can be viewed using FigTree software http://tree.bio.ed.ac.uk/software/figtree/

Two complete Alpha adintovirus-like contigs (NKLS02000104, NKLS02001728) were observed in assemblies of a PacBio-based WGS survey of bovine lung tissue. Sequences outside the inferred proviral inverted terminal repeats (ITRs) in the two sequences were highly diverse and mostly unidentifiable, but in a few reads the extra-proviral host sequences showed BLASTN similarity to genomic DNA sequences of various beetles, including *Tribolium castaneum* (a flour beetle that commonly infests cattle feed). Furthermore, the *Bos* lung-associated PolB sequences occupy phylogenetic clades comprised of insect-associated PolB sequences (Figure 3). These observations suggest that the two Alpha adintovirus sequences in the bovine datasets are insect-derived environmental contaminants, rather than mammal-tropic viruses. Similarly, several Beta adintovirus-like contigs (e.g., AANG04004209) found in a housecat oral swab sample show close phylogenetic affinity for adintovirus sequences observed in salmon WGS datasets. In another example, integrated adintoviruses found in a genomic dataset for olive trees (*Olea europaea*) showed insect-like sequences outside the inferred ITRs and showed phylogenetic affinity with PolB sequences from insect WGS datasets. Other adintovirus-like sequences found in plant datasets resembled adintovirus PolB sequences associated with nematode datasets. It thus appears that adintovirus sequences in some datasets are derived from environmental sources, as opposed to a productive infection of the organism that was the target of the sequencing effort.

Although there are examples of apparent environmental contamination, most adintovirus sequences form discrete clades that recapitulate the phylogeny of the host organisms that were the subjects of the WGS surveys. For example, a distinct clade of Alpha adintovirus PolB sequences was observed in datasets for multiple related species of venomous snakes. Several distinct clades of Beta adintoviruses were observed in datasets for amphibians and reptiles, including the well-populated clade that houses the exemplar *Terrapene* adintovirus. The exemplar *Mayetiola* adintovirus likewise occupies a clade exclusively populated by sequences found in insect WGS datasets.

TBLASTN searches against *Terrapene* box turtle Beta adintovirus PolB and Hexon protein sequences both yielded weak hits (E-value ~1e-05) for a locus on human chromosome 7. An adintovirus GasderminX sequence was also detected at the locus. Alignments to *Terrapene* adintovirus protein sequences were used to assign pseudogene annotations (Figure 4). The detected element is a highly disrupted endogenized Beta adintovirus. Homologous nucleotide sequences were detected in the genomes of primates, rodents, shrews, afrotherians, and xenarthrans but not in datasets for ungulates, carnivores, bats, marsupials, or prototherians. Endogenized adintovirus sequences observed in amphibian and reptile genomes do not share recognizable nucleotide similarity with placental mammal-endogenized adintovirus sequences. It is unclear whether a single adintovirus endogenization event affected an early placental mammal and the endogenized virus was then lost in non-shrew Laurasiatherians or whether multiple distinct endogenization events occurred in separate placental mammal lineages. Identification of extant examples of placental mammal adintoviruses could help resolve this question.

**Figure 4:**
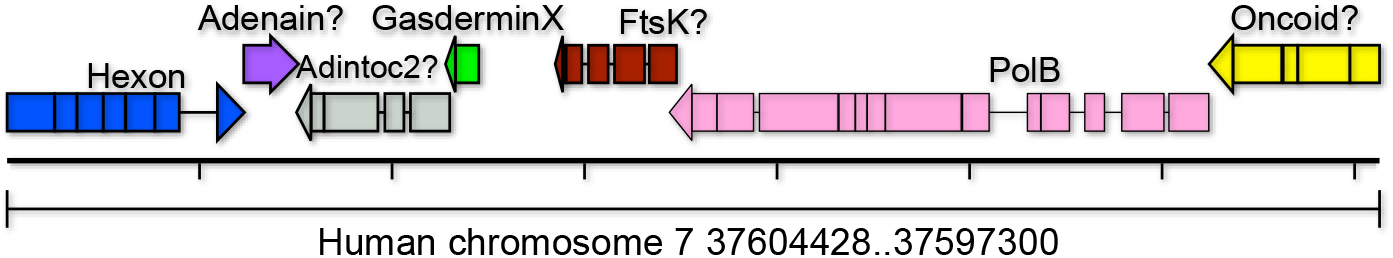
An endogenized Beta adintovirus relic found on human chromosome 7. Degraded pseudogenes interrupted by nonsense and frameshift mutations were reconstructed based on alignments to the protein sequences of *Terrapene* box turtle adintovirus. Question marks indicate that the reconstructed gene does not yield hits in BLAST searches of GenBank’s viruses taxon. Tentative gene assignments are based on synteny with the *Terrapene* adintovirus. The reconstructed Hexon, GasderminX, and PolB protein sequences yield DELTA-BLAST hits with E-values of 1e-21, 4e-05, and 3e-25, respectively. Figure supplement 1: annotated GenBank-format nucleotide map of the human chromosome 7 endogenized adintovirus depicted graphically in Figure 4.

### Viruses with adinto-like genes in non-animal eukaryote datasets

Eukaryotic viruses with midsize (10-50 kb) linear dsDNA genomes show a remarkable degree of genomic modularity (Koonin, Dolja et al. 2015, Yutin, Kapitonov et al. 2015, Yutin, Shevchenko et al. 2015). The apparently promiscuous horizontal gene transfer and lack of any single defining gene for these viruses makes the group taxonomically challenging. We propose the collective acronym MELD (midsize eukaryotic linear dsDNA) virus for the dizzyingly polyphyletic category. The name, which would encompass adenoviruses and adintoviruses, is intended to fill a gap between other operationally defined umbrella groups, such as CRESS viruses, small DNA tumor viruses, nucleocytoplasmic large DNA viruses, and megaviruses.

Datasets for *Blastocystis hominis* (a diatom-related unicellular eukaryote that commonly inhabits the human gut) contain MELD virus sequences that unite Alpha adintovirus-like PolB and integrase genes with inferred virion proteins whose primary sequences are not recognizably similar to known virion proteins (Figure 5). Gene identities for the *Blastocystis* virus were inferred based on HHpred results. Comparable MELD viruses were confirmed in rumen metagenomic datasets for sheep (Yutin, Kapitonov et al. 2015) and cattle, as well as in WGS datasets for green algae and fungi. In phylogenetic analyses, the PolB sequences of these viruses occupy long branches that are distant from animal-associated PolB clades (Figure 3).

**Figure 5:**
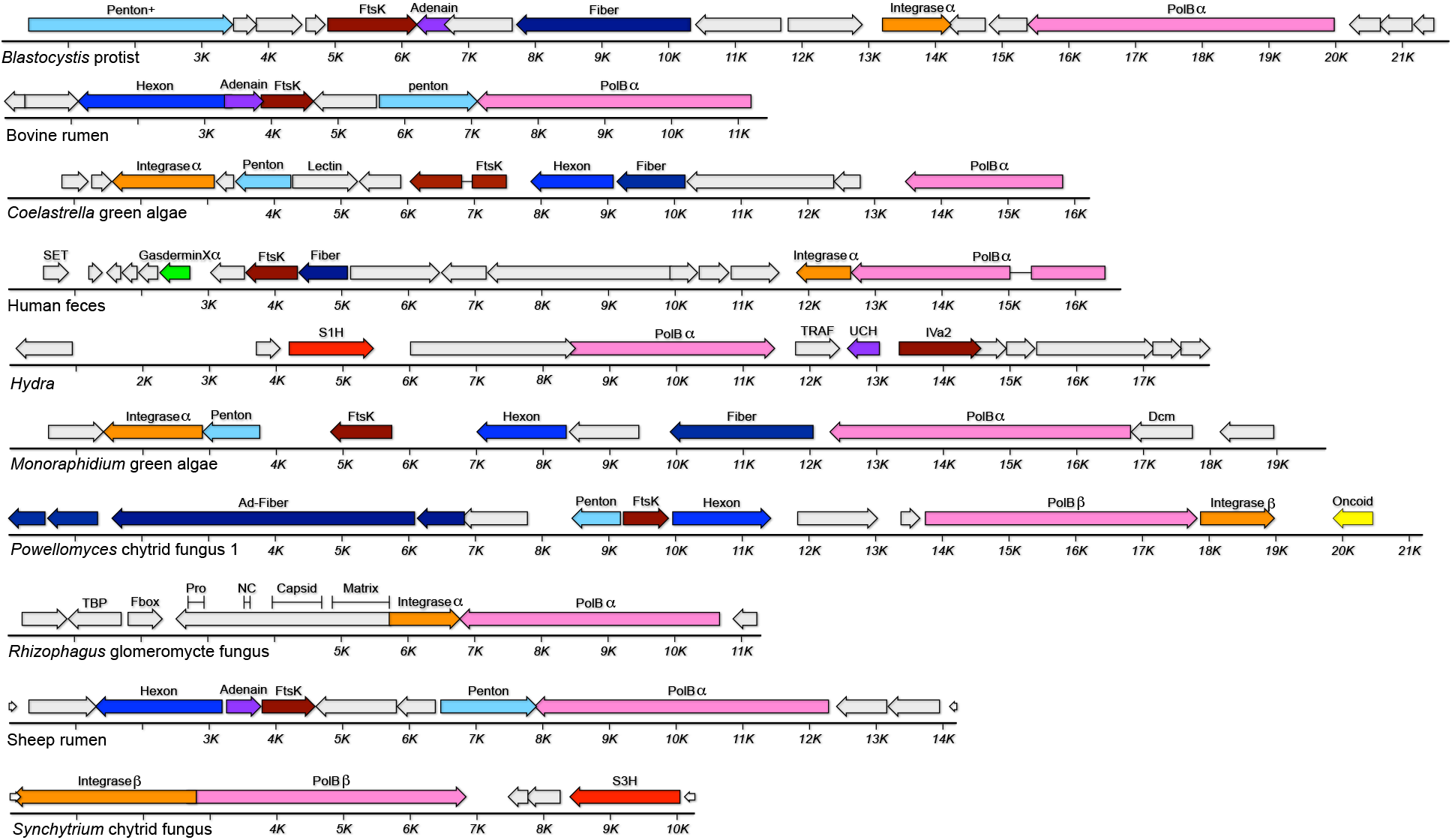
MELD viruses (and related elements) with adintovirus-like PolB genes. Greek letters indicate genes similar to adintoviruses in BLASTP or DELTA-BLAST searches. “Ad-” indicates similarity to adenovirus sequences. Abbreviations: Fiber, predicted structural or primary sequence similarity to bacteriophage tail fibers or coiled-coil proteins; Lectin, predicted structural similarity to galactose-binding domains; Dcm, predicted structural similarity to cytosine DNA methyltransferases; TBP, similar to TATA binding proteins; Matrix/Capsid/NC/Pro, similarities to retroviral Gag and retropepsin; S3H, poxvirus D5-like superfamily 3 helicase.

Contigs encoding Alpha adintovirus-like PolB and integrase genes were found in metagenomics datasets for bioreactor-cultured human feces, human urine samples, and human oral swab samples (Santiago-Rodriguez, Ly et al. 2015). This group of closely related sequences was only detected in datasets from a single laboratory and not in other human metagenomics surveys. Divergent variants of predicted proteins from the feces-associated virus were found in contigs from datasets for *Cyanophora paradoxa*, a species of glaucophyte algae (e.g., QPMI01000557), suggesting that the human feces-associated adintovirus-like sequences were derived from an environmental source.

Six MELD virus genomes assembled from a single *Powellomyces* SRA dataset unite sequences resembling adenovirus vertex fiber proteins with either a Beta adintovirus-like PolB (Figure 5) or a surprising variety of non-PolB DNA replicases (Figure 6). MELD virus genomes encoding genes similar to Alpha adintovirus virion proteins (E-values ~1e-7 to 1e-21) were assembled from datasets for *Capsaspora owczarzaki, Monosiga brevicollis,* and *Trichoplax H2* (unicellular eukaryotes that are thought to be closely related to animals). Aside from the abovementioned insect- and nematode-associated adintovirus sequences found in datasets for plants, adintovirus-like virion protein sequences were not detected in datasets for other non-animal eukaryotes. *Capsaspora* MELD virus 1 and the *Trichoplax* MELD virus both encode superfamily 1 helicase (S1H) genes instead of a PolB gene. Various megaviruses and bacteriophages encode similar S1H genes, as does a MELD virus observed in *Physarum polycephalum* slime mold and in *Powellomyces* MELD virus 4. Full-length S1H replicase genes of this class were not detected in animal WGS datasets, with the exception of seemingly endogenized degraded virus-like contigs in datasets for several coral and jellyfish species and a helitron-like element found in *Branchiostoma* lancelets (e.g., RDEB01009762, ABEP02037959).

**Figure 6:**
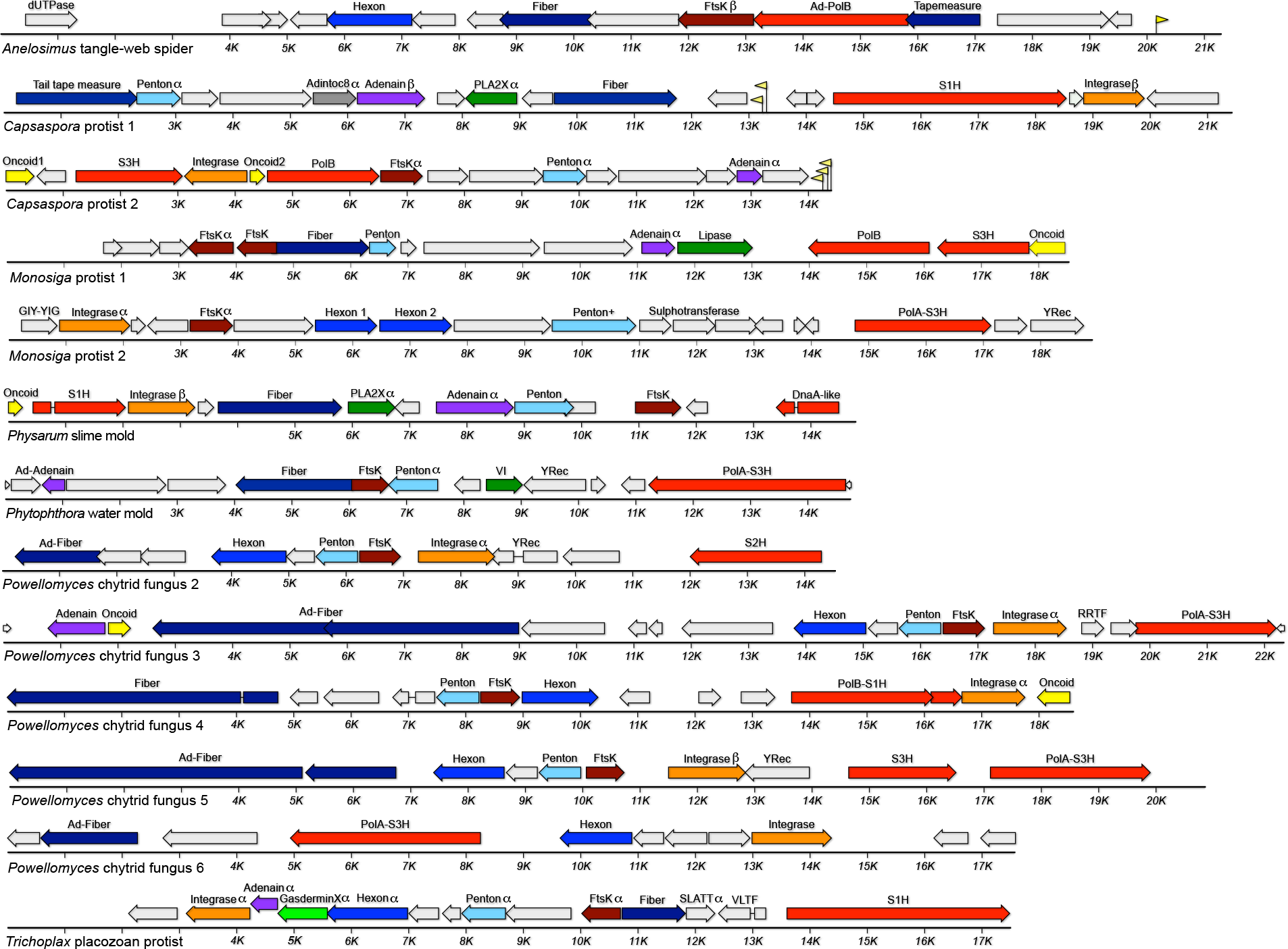
MELD viruses with other replicases. Greek letters indicate genes with sequences similar to Alpha or Beta adintoviruses in BLASTP searches. Sequences similar to adenoviruses are marked with “Ad-.” Yellow flags represent predicted tRNA genes. Abbreviations: dUTPase, similar to poxvirus deoxy-UTP diphosphatases; Fiber, similarity to bacteriophage tail fibers or other coiled-coil proteins; Tapemeasure, similarity to phage tail tape measure proteins; S1H, RecD/Pif1-like superfamily 1 helicase; YRec, homolog of phage tyrosine recombinases; TRAF, predicted structural similarity to TNF receptor associated factor 3; UCH, predicted structural similarity to ubiquitin C-terminal hydrolase cysteine proteases; IVa2, sequence similarity to adenovirus pIVa2 viral genome-packaging ATPases; DnaA-like, sequence distantly similar to DnaA and DnaB-like helicases; VI, similar to adenovirus virion core protein six; PolA, DNA polymerase family A (Pfam:00476); S3H superfamily 3 helicase similar to those observed in virophages and megaviruses; S2H, superfamily 2 helicase similar to DEAD-box helicase of Yellowstone Lake virophage 7 (YP_009177696); TRAF UCH TBP Fbox, homologs of host proteins with these gene symbols; SLATT, homolog of host SMODS and SLOG-associating 2TM effector domain proteins; VLTF, homolog of mimivirus VLTF3-like transcription factor. See main text for information about other gene names.

In WGS searches for sequences resembling human adenovirus type 5 PolB, we did not detect any contigs resembling full-length viruses in non-animal datasets. The searches did reveal the complete ITR-bounded genome of a typical mastadenovirus in a dataset for *Dipodomys ordii* (a type of kangaroo rat) as well as apparently complete MELD viruses in datasets for *Hydra oligactis* (brown hydra) and *Anelosimus studiosus* (a type of tangle-web spider). Like known adenoviruses, the *Hydra* and *Anelosimus* MELD viruses do not encode integrase genes and their PolB genes do not encode detectable OTU domains.

### PolB^+^ parvoviruses

BLASTP searches using Alpha adintovirus PolB sequences return high-likelihood matches (E-values <1e-80) for the PolB genes of an emerging group of bipartite parvoviruses referred to as bidnaviruses (Krupovic and Koonin 2014)(Figure 1). Like adintoviruses, bidnavirus PolB genes encode an N-terminal OTU domain. We searched assemblies of SRA datasets of interest for additional examples of bidnavirus genomes. PolB^+^ contigs were detected in datasets for the gut contents of African termites (*Cubitermes ugandensis*), dog (*Canis lupus familiaris*) feces, the silk glands of a false wolf spider (*Tengella perfuga*), and Tasmanian devil (*Sarcophilus harrisii*) feces (Figure 7). The dog feces PolB sequence is 53% similar to the “structural protein” of Fresh Meadows “densovirus” 3 previously detected in mouse (*Mus musculus*) feces (AWB14611)(Williams, Che et al. 2018).

**Figure 7:**
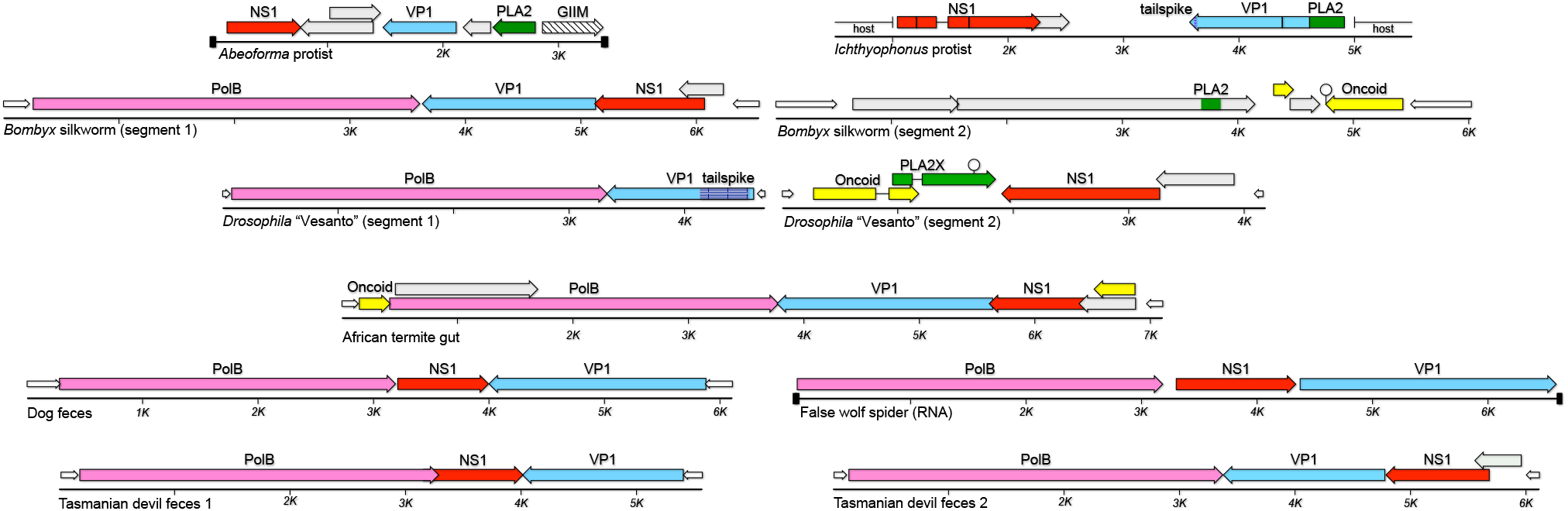
Bidnaparvovirus and non-animal parvovirus genome maps. Abbreviations: GIIM, similarity to group II intron maturases; tailspike, similarity to bacteriophage short tail fibers.

In previously reported bidnaviruses, the termini of each of the two genome segments have matching nucleotide sequences. To search for second segments, we probed the assemblies for examples of other contigs with termini similar to the ITRs of the initially observed bidnavirus contigs. The datasets were also searched for contigs with sequences similar to previously reported bidnavirus proteins. Second segments were not detected, suggesting that the five new bidnaviruses may be monopartite. We suggest that the apparently monopartite viruses could still be referred to as bidnaviruses (or, more specifically, bidnaparvoviruses) but with the “bidna” moniker connoting the presence of two types of DNA replicase genes, as opposed to the original connotation of a virus with two genomic DNA segments.

Searches for examples of parvovirus NS1-like sequences did not reveal clear examples outside of multicellular animal datasets. A marginal exception was a group of sequences found in datasets for *Abeoforma whisleri* and *Ichthyophonus hoferi*, two unicellular eukaryotes that are thought to be closely related to multicellular animals. The observations suggest an early-animal origin for parvoviruses that involved acquisition of genes from Alpha adintoviruses.

## Discussion

We have identified a coherent family-like grouping of animal viruses that we call adintoviruses, connoting their hallmark adenovirus-like virion protein genes and retrovirus-like integrase genes. Adintovirus sequences are detectable either as apparently free linear DNA molecules or as endogenized integrants in WGS datasets representing all eumetazoan phyla. Although sequences resembling the PolB proteins of Alpha and Beta adintoviruses can also be found in datasets for non-animal eukaryotes, the sequences of adintovirus virion proteins appear to be restricted to animals.

We imagine that the related Alpha and Beta adintovirus-like lineages might have infected early eukaryotes and the two lineages gradually co-evolved with major divisions of eukaryotes, including multicellular animals. In this model, the sequences of the virion protein genes presumably evolved more rapidly than the conserved catalytic core of PolB, resulting in distinctive mutually unrecognizable virion protein sequences specific to each major division of eukaryotes. The model suggests that adenoviruses could be thought of as a related sister lineage that also arose in or before the first animals. Although adenoviruses are currently only known to infect vertebrates, the idea that the lineage long predates the emergence of vertebrates is consistent with our identification of a distantly adenovirus-like sequence in a spider dataset (Figure 6).

In non-animal eukaryote datasets, adintovirus-like PolB sequences can be found in a wide range of sequence contexts, ranging from elements with no obvious virion proteins to the genomes of megaviruses. Conversely, it appears that the adintovirus-like PolB can readily be replaced with other types of DNA replicase genes (Figure 6). This presumably reflects the previously proposed rampant horizontal gene transfer among virus lineages that infect unicellular eukaryotes. Although similar horizontal gene transfer events appear to have occurred between various animal-tropic virus families, including adintoviruses and parvoviruses (Figure 7), adomaviruses and polyomaviruses (Mizutani, Sayama et al. 2011, Dill, Camus et al. 2018) https://www.biorxiv.org/content/10.1101/341131v2 and papillomaviruses and polyomaviruses (Woolford, Rector et al. 2007), each of these cases appears to represent single ancient event. It may be that the evolution of distinct tissues and organs or the development of cell-mediated immunity in multicellular animals placed limits on the likelihood that different virus lineages can co-infect a single cell and productively recombine. From this view, the distinctive gene combinations seen in adintoviruses and adenoviruses might simply be bottlenecked examples of the much larger range of gene combinations observed in MELD viruses of unicellular eukaryotes (Yutin, Kapitonov et al. 2015, Yutin, Shevchenko et al. 2015).

It has generally been assumed that the functionally similar oncogenes found in adenoviruses, papillomaviruses, parvoviruses, and polyomaviruses arose through convergent evolution or through horizontal gene transfer between virus families (de Souza, Iyer et al. 2010). Although small DNA tumor virus oncogenes show low overall sequence similarity, they can be roughly defined based on the presence of short linear motifs. Many adintoviruses encode candidate “Oncoid” proteins with these motifs. Bombyx silkworm bidnaparvovirus NS3, which we have designated as a candidate Oncoid (Figure 7) has previously been shown to be similar to a baculovirus protein of unknown function (Krupovic and Koonin 2014). We note that many of the proposed baculovirus homologs (e.g., YP_009506034) share potential zinc-coordinating cysteine residues as well as a C-terminal LXCXE/CK2 site, qualifying the baculovirus proteins as candidate Oncoids as well. Surprisingly, candidate Oncoids were also observed in MELD viruses of unicellular eukaryotes (Figure 6). The predicted Oncoid2 gene of *Capsaspora* protist MELD virus 2 detects polyomavirus Large T oncogenes in DELTA-BLAST searches (E-value 4e-16). It is interesting to imagine that oncogenes in a broad range of animal DNA viruses might share an ancestry that pre-dates the emergence of multicellular animals.

Adintoviruses encode a number of accessory genes that appear to be homologs of membrane-active proteins found in animal venom. These include bee and snake venom PLA2 and melittin, as well as a spider venom protein called cupiennin. Interestingly, venom PLA2 and melittin (which shows similarity to adenovirus pX in HHpred searches) act in concert (Vogt, Patzer et al. 1970), suggesting the speculative hypothesis that these venom genes might have arisen from a captured viral PLA2X-like gene.

In unpublished work, our group used a standard baculovirus-based expression system (ThermoFisher) to generate a virus-like particle (VLP) vaccine against BK polyomavirus (BKV)(Peretti, Geoghegan et al. 2018). The project provided an inadvertent natural experiment. Recombinant baculoviruses were generated in Sf9 cells and bulk protein expression was performed using the *Trichoplusia ni* cell line High Five. BKV VLPs were purified according to previously reported methods (Cardone, Moyer et al. 2014) involving ultracentrifugation through density gradients, nuclease digestion, and size exclusion chromatography. Deep sequencing of DNA extracted from the purified VLP preparation shows high-depth coverage of *Spodoptera* adintovirus genomes alongside incomplete patchy coverage of endogenized *Trichoplusia*-specific homologs of the two *Spodoptera* viruses (Supplemental File 1). It appears that Sf9-derived adintoviruses infected the High Five cells and this led to the production of adintovirus virions that co-purified with the recombinant BKV VLPs. The results suggest that standard insect cell cultures could serve as a laboratory model for productive adintovirus infection.

A Beta adintovirus was detected in transcriptomic and WGS datasets for Mexican blind tetra cavefish (*Astyanax mexicanus*). Adintovirus transcripts were most abundant in head, kidney, and intestine samples and least abundant in muscle and whole embryo samples (Supplemental Table 1). Analysis of the WGS dataset showed that adintovirus DNA reads outnumbered reads for a single copy host gene (gamma tubulin, NW_019172896) by a factor of 25. At both an RNA and DNA level the *Astyanax* sequence showed a high degree of uniformity, suggesting a clonal infection. In contrast, pet store samples of a different species of tetra, *Gymnocorymbus ternetzi* (SRR2040422), showed such a complex range of adintovirus sequence variants that assembly of contigs representing complete viral genomes was challenging. These observations suggest that tetras might serve as a tractable laboratory model for adintovirus infection.

Adintoviruses have a number of features that could make them useful as recombinant gene transfer vectors. Their genome size is substantially larger than commonly used retroviral and parvoviral vectors. In contrast to adenovirus- and baculovirus-based vector systems, adintovirus genomes are small enough to be manipulated entirely in the setting of standard plasmids. An intriguing feature of the adintovirus integrase gene is the presence of a predicted chromodomain that, in LTR retrotransposons, is believed to influence integration site specificity (Kordis 2005). This could theoretically offer an advantage over retroviral vectors, which show little integration site specificity. Another potential practical use for adintoviruses might be as biocontrol agents for pest organisms, such as *Mayetiola destructor* barley midges or chytrid fungi that parasitize amphibians.

An important implication of this study is that there may be additional unappreciated families of animal viruses hiding in plain sight in sequence databases. Adintoviruses may have been relatively easy quarry because they are able to integrate into host genomes, such that they are detectable in WGS datasets of randomly sampled animals that did not happen to be suffering from an active infection. In contrast to the hundreds of adintovirus-like contigs detected in our initial WGS survey, focused searches for adenoviruses (which do not encode integrases) detected only a single complete adenovirus genome. For future discovery efforts, it will be important to develop higher throughput methods using sensitive structure-guided searches to identify divergent new examples of viral hallmark genes in sequence datasets representing many individuals, including subjects suffering from disease. The key goal will be to understand which combinations of genes tend to co-occupy single contigs. Recently reported bioinformatics pipelines, such as Cenote-Taker (Tisza, Pastrana et al. 2019) and Mash Screen (Ondov, Starrett et al. 2019), should be useful for these purposes. Deposition of annotated viral genome sequences into publicly searchable databases will be critical for further expanding our understanding of the eukaryotic virome.

## Materials and Methods

### Detection and analysis of viral sequences

Adomavirus LO8 (Adenain) sequences were initially used for TBLASTN searches of the NCBI TSA and WGS databases. The relationship between adomavirus and adintovirus virion proteins is the subject of a separate manuscript (BioRxiv 341131v2). The Adenain sequences of *Nephila* orb-weaver spider contig (GFKT014647032) or a *Parasteatoda* spider contig (AOMJ02256338) were arbitrarily chosen for further TBLASTN searches of eukaryotic datasets in TSA and WGS databases. Adenain-bearing contigs 4-50kb in length were further searched (using CLC Genomics Workbench) for BLASTP-detectable PolB homologs. Contigs were inspected for the presence of nearly overlapping arrays of large (>100 AA) open reading frames. Contigs with inverted repeats flanking the ORF cluster were favored, but this was not a strict sorting criterion.

Selected contigs of interest were initially annotated using DELTA-BLAST searches of GenBank nr or HHpred analyses of single or aligned protein sequences against PDB_mmCIF70, COG_KOG, Pfam-A, and NCBI_CD databases (Altschul, Madden et al. 1997, Altschul, Wootton et al. 2005, Soding 2005, Hildebrand, Remmert et al. 2009, Gerlt, Bouvier et al. 2015, Meier and Soding 2015, Zimmermann, Stephens et al. 2017). Protein sequences were extracted from the contigs using getORF http://bioinfo.nhri.org.tw/cgi-bin/emboss/getorf (Rice, Longden et al. 2000). Extracted protein sequences were clustered using EFI-EST https://efi.igb.illinois.edu/efi-est/ (Gerlt, Bouvier et al. 2015, Zallot, Oberg et al. 2018) and displayed using Cytoscape v3.7.1 (Shannon, Markiel et al. 2003). Multiple sequence alignments were constructed using MAFFT https://toolkit.tuebingen.mpg.de/#/tools/mafft. Contigs were annotated using Cenote-Taker (Tisza, Pastrana et al. 2019) with an iteratively refined library of conserved adintovirus protein sequences. Compiled protein sequences are provided as a zipped set of fasta-format text files in Figure 1 Figure supplement 2. Maps were drawn using MacVector 17 software. Phylogenetic analyses were performed using MAFFT 7 https://mafft.cbrc.jp/alignment/server/ (Kuraku, Zmasek et al. 2013, Katoh, Rozewicki et al. 2019) and displayed using FigTree 1.4.4 http://tree.bio.ed.ac.uk/software/figtree/.

Selected contigs for which SRA datasets were available were subjected to reference-guided re-assembly using Megahit 1.2.9 (Li, Liu et al. 2015, Li, Luo et al. 2016) and/or the map reads to reference function of CLC Genomics Workbench. Annotated maps were submitted to GenBank as third party annotation assemblies (TPA_asm). Graphical examples of the annotation process are depicted in Figure 2 Figure supplement 2.

## Supporting information

Fig1 Supp1 PolB Cytoscape

Fig1 Supp2 Protein compilations

Fig1 Supp3 Hexon Penton networks

Fig2 Supp1 Accession numbers

Fig2 Supp2 Annotation examples

Fig2 Supp3 Additional Adintos

Fig2 Supp4 Nucleotide maps

Fig3 Supp1&2 WGS PolB trees

Fig4 Supp1 Human Adinto nucleotide

Supp File1 VLP reads

Supp Table 1 Astyanax expression

## Data Availability

GenBank accession numbers for sequences deposited in association with this study are: BK010888 BK010889 BK010890 BK010893 BK010894 BK010998 BK010999 BK011000 BK011001 BK011002 BK011003 BK011004 BK011005 BK011006 BK011007 BK011008 BK011009 BK011010 BK011011 BK011022 BK011023 BK011024 BK011025 BK011026 BK012042 BK012043 BK012044 BK012045 BK012046 BK012047 BK012048 BK012049 BK012050 BK012051 BK012052 BK012053 BK012054 BK012055 BK012056 BK012057 BK012058 BK012059 BK012060 BK012061 BK012062 BK012063 BK012064 BK012084 BK012085 BK012086.

## Acknowledgments

The authors are indebted to Eugene Koonin and Natalya Yutin for their generous guidance and for the spirited discussions that inspired us to pursue this study. We thank Karl Münger and Jim Pipas for their extensive advice about oncogene sequence motifs and Chris Bellas for useful discussions and for critical evaluation of the manuscript. We are also grateful to Patrick McTamney for guiding the production of BK polyomavirus VLP stocks.

## References

Altschul, S. F., T. L. Madden, A. A. Schaffer, J. Zhang, Z. Zhang, W. Miller and D. J. Lipman (1997). “Gapped BLAST and PSI-BLAST: a new generation of protein database search programs.” Nucleic Acids Res 25(17): 3389–3402.

Altschul, S. F., J. C. Wootton, E. M. Gertz, R. Agarwala, A. Morgulis, A. A. Schaffer and Y. K. Yu (2005). “Protein database searches using compositionally adjusted substitution matrices.” FEBS J 272(20): 5101–5109.

Boratyn, G. M., A. A. Schaffer, R. Agarwala, S. F. Altschul, D. J. Lipman and T. L. Madden (2012). “Domain enhanced lookup time accelerated BLAST.” Biol Direct 7: 12.

Buck, C. B., K. Van Doorslaer, A. Peretti, E. M. Geoghegan, M. J. Tisza, P. An, J. P. Katz, J. M. Pipas, A. A. McBride, A. C. Camus, A. J. McDermott, J. A. Dill, E. Delwart, T. F. Ng, K. Farkas, C. Austin, S. Kraberger, W. Davison, D. V. Pastrana and A. Varsani (2016). “The Ancient Evolutionary History of Polyomaviruses.” PLoS Pathog 12(4): e1005574.

Cardone, G., A. L. Moyer, N. Cheng, C. D. Thompson, I. Dvoretzky, D. R. Lowy, J. T. Schiller, A. C. Steven, C. B. Buck and B. L. Trus (2014). “Maturation of the human papillomavirus 16 capsid.” MBio 5(4): e01104–01114.

de Souza, R. F., L. M. Iyer and L. Aravind (2010). “Diversity and evolution of chromatin proteins encoded by DNA viruses.” Biochim Biophys Acta 1799(3-4): 302–318.

Dill, J. A., A. C. Camus, J. H. Leary and T. F. F. Ng (2018). “Microscopic and Molecular Evidence of the First Elasmobranch Adomavirus, the Cause of Skin Disease in a Giant Guitarfish, Rhynchobatus djiddensis.” MBio 9(3).

Dubois, H., F. Sorgeloos, S. T. Sarvestani, L. Martens, Y. Saeys, J. M. Mackenzie, M. Lamkanfi, G. van Loo, I. Goodfellow and A. Wullaert (2019). “Nlrp3 inflammasome activation and Gasdermin D-driven pyroptosis are immunopathogenic upon gastrointestinal norovirus infection.” PLoS Pathog 15(4): e1007709.

Duponchel, S. and M. G. Fischer (2019). “Viva lavidaviruses! Five features of virophages that parasitize giant DNA viruses.” PLoS Pathog 15(3): e1007592.

Fischer, M. G. and T. Hackl (2016). “Host genome integration and giant virus-induced reactivation of the virophage mavirus.” Nature 540(7632): 288–291.

Fu, L., B. Niu, Z. Zhu, S. Wu and W. Li (2012). “CD-HIT: accelerated for clustering the next-generation sequencing data.” Bioinformatics 28(23): 3150–3152.

Geisler, C. (2018). “A new approach for detecting adventitious viruses shows Sf-rhabdovirus-negative Sf-RVN cells are suitable for safe biologicals production.” BMC Biotechnol 18(1): 8.

Gerlt, J. A., J. T. Bouvier, D. B. Davidson, H. J. Imker, B. Sadkhin, D. R. Slater and K. L. Whalen (2015). “Enzyme Function Initiative-Enzyme Similarity Tool (EFI-EST): A web tool for generating protein sequence similarity networks.” Biochim Biophys Acta 1854(8): 1019–1037.

Gouw, M., S. Michael, H. Samano-Sanchez, M. Kumar, A. Zeke, B. Lang, B. Bely, L. B. Chemes, N. E. Davey, Z. Deng, F. Diella, C. M. Gurth, A. K. Huber, S. Kleinsorg, L. S. Schlegel, N. Palopoli, K. V. Roey, B. Altenberg, A. Remenyi, H. Dinkel and T. J. Gibson (2018). “The eukaryotic linear motif resource - 2018 update.” Nucleic Acids Res 46(D1): D428–D434.

Hildebrand, A., M. Remmert, A. Biegert and J. Soding (2009). “Fast and accurate automatic structure prediction with HHpred.” Proteins 77 Suppl 9: 128–132.

Huang, Y., B. Niu, Y. Gao, L. Fu and W. Li (2010). “CD-HIT Suite: a web server for clustering and comparing biological sequences.” Bioinformatics 26(5): 680–682.

Iyer, L. M., K. S. Makarova, E. V. Koonin and L. Aravind (2004). “Comparative genomics of the FtsK-HerA superfamily of pumping ATPases: implications for the origins of chromosome segregation, cell division and viral capsid packaging.” Nucleic Acids Res 32(17): 5260–5279.

Katoh, K., J. Rozewicki and K. D. Yamada (2019). “MAFFT online service: multiple sequence alignment, interactive sequence choice and visualization.” Brief Bioinform 20(4): 1160–1166.

Koonin, E. V., V. V. Dolja and M. Krupovic (2015). “Origins and evolution of viruses of eukaryotes: The ultimate modularity.” Virology 479-480: 2–25.

Koonin, E. V., M. Krupovic and N. Yutin (2015). “Evolution of double-stranded DNA viruses of eukaryotes: from bacteriophages to transposons to giant viruses.” Ann N Y Acad Sci 1341: 10–24.

Kordis, D. (2005). “A genomic perspective on the chromodomain-containing retrotransposons: Chromoviruses.” Gene 347(2): 161–173.

Krupovic, M., D. H. Bamford and E. V. Koonin (2014). “Conservation of major and minor jelly-roll capsid proteins in Polinton (Maverick) transposons suggests that they are bona fide viruses.” Biol Direct 9: 6.

Krupovic, M. and E. V. Koonin (2014). “Evolution of eukaryotic single-stranded DNA viruses of the Bidnaviridae family from genes of four other groups of widely different viruses.” Sci Rep 4: 5347.

Krupovic, M. and E. V. Koonin (2015). “Polintons: a hotbed of eukaryotic virus, transposon and plasmid evolution.” Nat Rev Microbiol 13(2): 105–115.

Kuraku, S., C. M. Zmasek, O. Nishimura and K. Katoh (2013). “aLeaves facilitates on-demand exploration of metazoan gene family trees on MAFFT sequence alignment server with enhanced interactivity.” Nucleic Acids Res 41(Web Server issue): W22–28.

Li, D., C. M. Liu, R. Luo, K. Sadakane and T. W. Lam (2015). “MEGAHIT: an ultra-fast single-node solution for large and complex metagenomics assembly via succinct de Bruijn graph.” Bioinformatics 31(10): 1674–1676.

Li, D., R. Luo, C. M. Liu, C. M. Leung, H. F. Ting, K. Sadakane, H. Yamashita and T. W. Lam (2016). “MEGAHIT v1.0: A fast and scalable metagenome assembler driven by advanced methodologies and community practices.” Methods 102: 3–11.

Li, W. and A. Godzik (2006). “Cd-hit: a fast program for clustering and comparing large sets of protein or nucleotide sequences.” Bioinformatics 22(13): 1658–1659.

Meier, A. and J. Soding (2015). “Automatic Prediction of Protein 3D Structures by Probabilistic Multi-template Homology Modeling.” PLoS Comput Biol 11(10): e1004343.

Mizutani, T., Y. Sayama, A. Nakanishi, H. Ochiai, K. Sakai, K. Wakabayashi, N. Tanaka, E. Miura, M. Oba, I. Kurane, M. Saijo, S. Morikawa and S. Ono (2011). “Novel DNA virus isolated from samples showing endothelial cell necrosis in the Japanese eel, Anguilla japonica.” Virology 412(1): 179–187.

Moriyama, T., K. Terasawa, M. Fujiwara and N. Sato (2008). “Purification and characterization of organellar DNA polymerases in the red alga Cyanidioschyzon merolae.” FEBS J 275(11): 2899–2918.

Ondov, B. D., G. J. Starrett, A. Sappington, A. Kostic, S. Koren, C. B. Buck and A. M. Phillippy (2019). “Mash Screen: High-throughput sequence containment estimation for genome discovery.” bioRxiv: 557314.

Peretti, A., E. M. Geoghegan, D. V. Pastrana, S. Smola, P. Feld, M. Sauter, S. Lohse, M. Ramesh, E. S. Lim, D. Wang, C. Borgogna, P. C. FitzGerald, V. Bliskovsky, G. J. Starrett, E. K. Law, R. S. Harris, J. K. Killian, J. Zhu, M. Pineda, P. S. Meltzer, R. Boldorini, M. Gariglio and C. B. Buck (2018). “Characterization of BK Polyomaviruses from Kidney Transplant Recipients Suggests a Role for APOBEC3 in Driving In-Host Virus Evolution.” Cell Host Microbe 23(5): 628–635 e627.

Pipas, J. M. (1992). “Common and unique features of T antigens encoded by the polyomavirus group.” J Virol 66(7): 3979–3985.

Pipas, J. M. (2019). “DNA Tumor Viruses and Their Contributions to Molecular Biology.” J Virol 93(9).

Rice, P., I. Longden and A. Bleasby (2000). “EMBOSS: the European Molecular Biology Open Software Suite.” Trends Genet 16(6): 276–277.

Santiago-Rodriguez, T. M., M. Ly, M. C. Daigneault, I. H. Brown, J. A. McDonald, N. Bonilla, E. A. Vercoe and D. T. Pride (2015). “Chemostat culture systems support diverse bacteriophage communities from human feces.” Microbiome 3: 58.

Shannon, P., A. Markiel, O. Ozier, N. S. Baliga, J. T. Wang, D. Ramage, N. Amin, B. Schwikowski and T. Ideker (2003). “Cytoscape: a software environment for integrated models of biomolecular interaction networks.” Genome Res 13(11): 2498–2504.

Soding, J. (2005). “Protein homology detection by HMM-HMM comparison.” Bioinformatics 21(7): 951–960.

Tisza, M. J., D. V. Pastrana, N. L. Welch, B. Stewart, A. Peretti, G. J. Starrett, Y.-Y. S. Pang, A. Varsani, S. R. Krishnamurthy, P. A. Pesavento, D. H. McDermott, P. M. Murphy, J. L. Whited, B. Miller, J. M. Brenchley, S. P. Rosshart, B. Rehermann, J. Doorbar, B. A. Ta’ala, O. Pletnikova, J. Troncoso, S. M. Resnick, A. M. Segall and C. B. Buck (2019). “Discovery of several thousand highly diverse circular DNA viruses.” 555375.

Vogt, W., P. Patzer, L. Lege, H. D. Oldigs and G. Wille (1970). “Synergism between phospholipase A and various peptides and SH-reagents in causing haemolysis.” Naunyn Schmiedebergs Arch Pharmakol 265(5): 442–454.

Williams, S. H., X. Che, J. A. Garcia, J. D. Klena, B. Lee, D. Muller, W. Ulrich, R. M. Corrigan, S. Nichol, K. Jain and W. I. Lipkin (2018). “Viral Diversity of House Mice in New York City.” MBio 9(2).

Woolford, L., A. Rector, M. Van Ranst, A. Ducki, M. D. Bennett, P. K. Nicholls, K. S. Warren, R. A. Swan, G. E. Wilcox and A. J. O’Hara (2007). “A novel virus detected in papillomas and carcinomas of the endangered western barred bandicoot (Perameles bougainville) exhibits genomic features of both the Papillomaviridae and Polyomaviridae.” J Virol 81(24): 13280–13290.

Yutin, N., V. V. Kapitonov and E. V. Koonin (2015). “A new family of hybrid virophages from an animal gut metagenome.” Biol Direct 10: 19.

Yutin, N., D. Raoult and E. V. Koonin (2013). “Virophages, polintons, and transpovirons: a complex evolutionary network of diverse selfish genetic elements with different reproduction strategies.” Virol J 10: 158.

Yutin, N., S. Shevchenko, V. Kapitonov, M. Krupovic and E. V. Koonin (2015). “A novel group of diverse Polinton-like viruses discovered by metagenome analysis.” BMC Biol 13: 95.

Zallot, R., N. O. Oberg and J. A. Gerlt (2018). “‘Democratized’ genomic enzymology web tools for functional assignment.” Curr Opin Chem Biol 47: 77–85.

Zimmermann, L., A. Stephens, S. Z. Nam, D. Rau, J. Kübler, M. Lozajic, F. Gabler, J. Söding, A. N. Lupas and V. Alva (2017). “A Completely Reimplemented MPI Bioinformatics Toolkit with a New HHpred Server at its Core.” J Mol Biol S0022-2836(17): 30587–30589.

