## Supplementary material for "Adintoviruses: An Animal-Tropic Family of Midsize Eukaryotic Linear dsDNA (MELD) Viruses": Fig1 Supp3 Hexon Penton networks

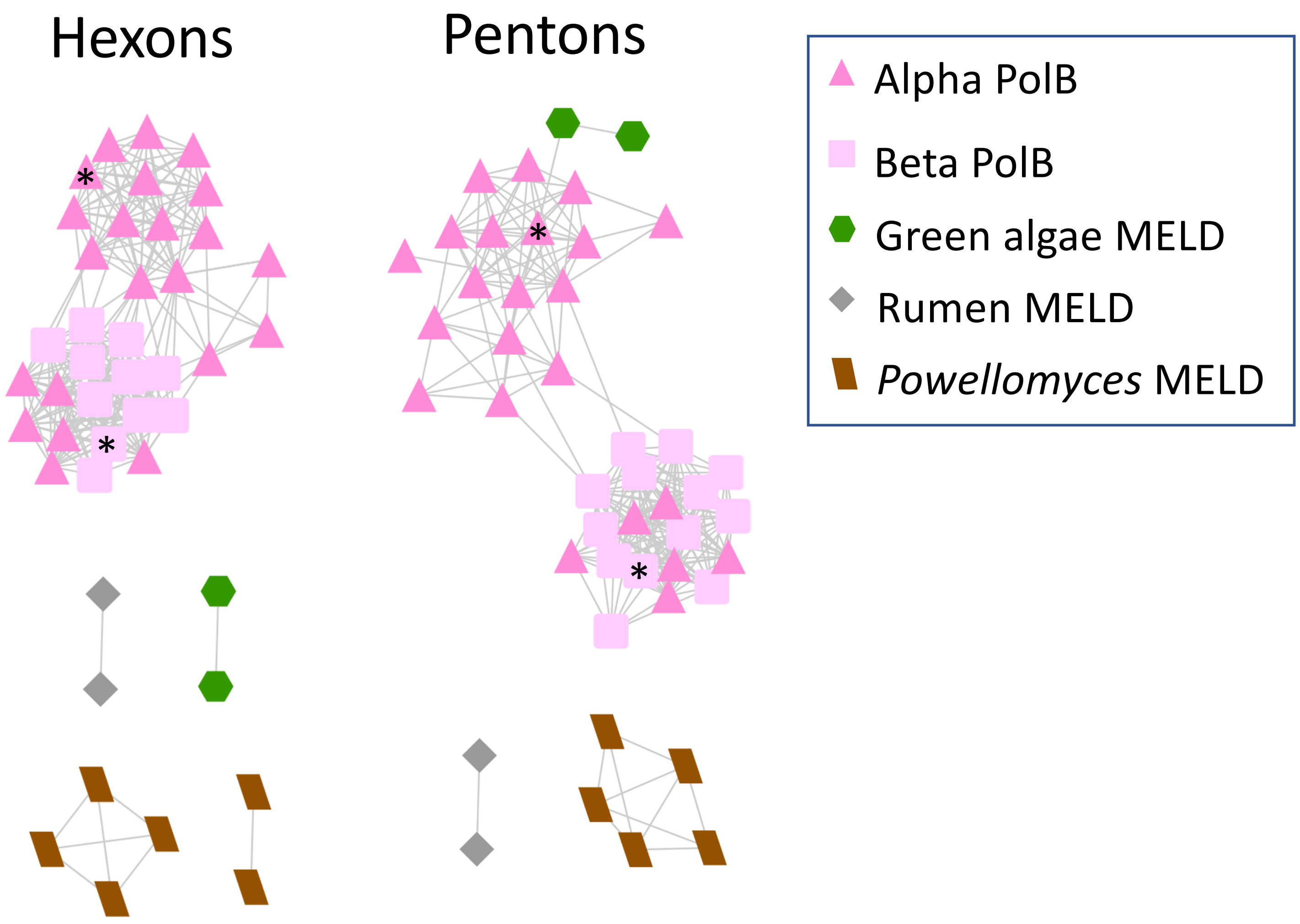


**Figure 2 Figure supplement 4: Sequence similarity network analysis for Hexon and Penton proteins on curated contigs.** Hexon and Penton protein sequences from curated adintovirus and MELD virus genomes (see Figure 2 Figure supplement 1 for accession numbers) were subjected to all-against-all BLASTP analysis with an E-value cutoff of 1e-5. Asterisks mark the *Mayetiola* (triangle) and *Terrapene* (square) exemplars. The figure shows that some genomes with Alpha PolB genes have Hexon and Penton proteins that cluster with Beta-type Hexon and Penton proteins. The figure also shows that non-animal MELD virus Hexon and Penton protein sequences generally cluster with one another and not with animal adintovirus sequences (see Results section "Viruses with adinto-like genes in non-animal eukaryote datasets").
