## Supplementary material for "Adintoviruses: An Animal-Tropic Family of Midsize Eukaryotic Linear dsDNA (MELD) Viruses": Fig2 Supp2 Annotation examples

### Annotation of Terrapene box turtle Beta adintovirus BK010890

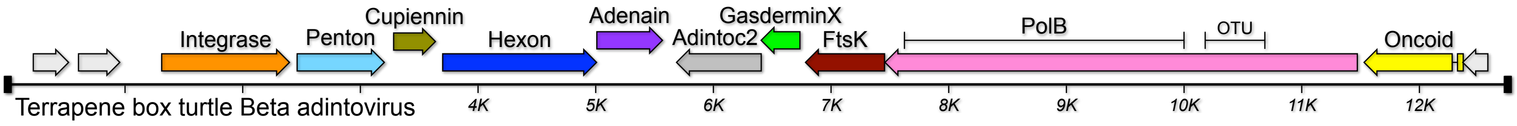


#### Integrase

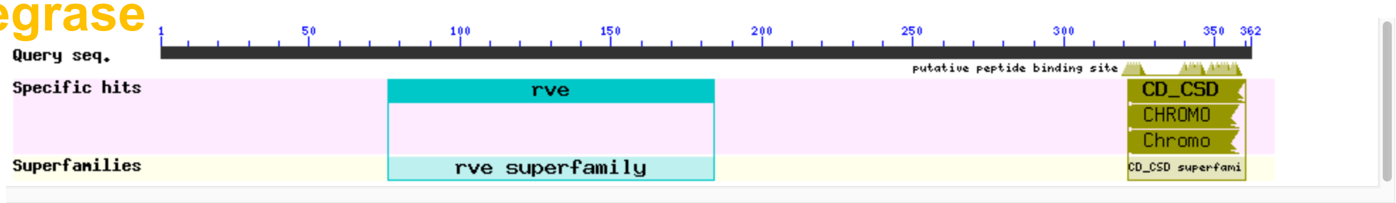


Search for similar domain architectures | Refine search

**List of domain hits**

| Name | Accession | Description | Interval | E-value |
|---|---|---|---|---|
| [+] rve | pfam00665 | Integrase core domain; Integrase mediates integration of a DNA copy of the viral genome into ... | 76-184 | 5.71e-18 |
| [+] CD_CSD | cd00024 | CHROMO (CHRromatin Organization Modifier) domains and chromo shadow domains; Members of this ... | 321-360 | 9.40e-09 |
| [+] CHROMO | smart00298 | Chromatin organization modifier domain; | 321-358 | 2.19e-06 |
| [+] Chromo | pfam00385 | Chromo (CHRromatin Organisation MOdifier) domain; | 321-358 | 2.52e-06 |

**Blast search parameters**

**Data Source:** Live blast search **RID =** W2GZC0Z5015

**User Options:** **Database:** DELTA_BLAST/cdd_delta **Low complexity filter:** no **Composition Based Adjustment:** yes **E-value threshold:** 0.05 **Maximum number of hits:** 500

#### Penton

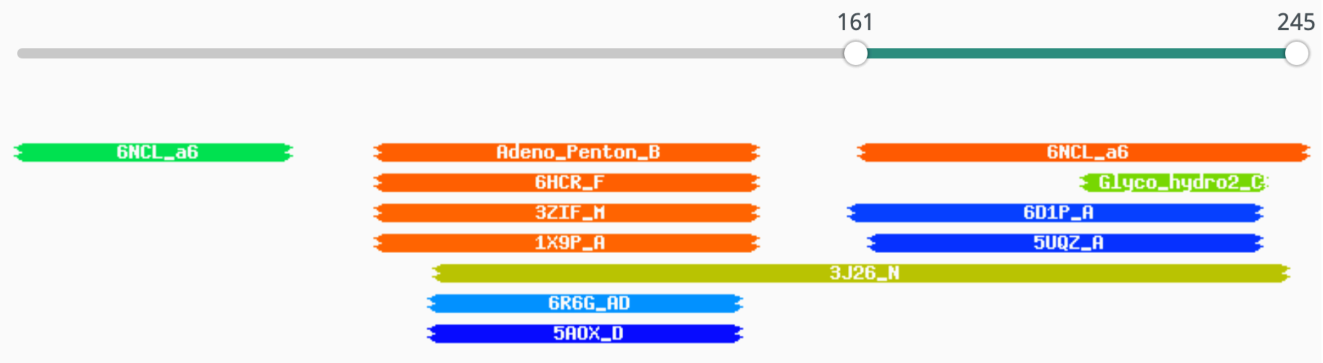


**PF01686.17 ; Adeno_Penton_B ; Adenovirus penton base protein**

Probability: 91.79%, E-value: 0.37, Score: 45.32, Aligned cols: 69, Identities: 19%, Similarity: 0.366,

```
Q ss_pred                  ceEEEEeCCcccCCHHHHHHHHHHhhccC------------------CCCCceeEEEeCCCC--------------EEEE
Q Q_TerrapenePen    70   TTWNFILRRGYYSTIPELLEHMNSNVARH------------------PGPPEVVMNYDPVGR--------------KVRL   117 (246)
Q Consensus        70   ~~~~~~ip~G~Y~~i~dL~~~ln~~l~~~------------------~~~~~~~~~~~~~~~~n~~~~--------------k~~i   117 (246)
                           +...++||+|.|+ ..++++.||+..-++                  -...+|.+.+|+.++              -+.+
T Consensus       124   eW~~ltIPEGNys-~~~~IDLlNnaIve~YL~~GRQngV~esDIGVKFDTRnF~Lg~DP~T~LVtPG~YtykafHPDIvL   202 (450)
T PF01686.17      124   EWVQITIPEGNYT-GSELIDQLNNGILNNYLEVGRQKGVEIEDIGVKFDTRDFSLGYDPETGLITPGKYTYKAFHPDIIL   202 (450)
T ss_pred                  EEEEEEecCCCCc-HHHHHHHHHHHHHHHHHHHHHHhCCcHHHceeeeeccccccCcCCCCCcccCccceecccCCCEEE


Q ss_pred                  EcCCcEEEEecCchHHHHhCCcCC
Q Q_TerrapenePen   118   KSTDFMYVFSTGGELANILGLGHK   141 (246)
Q Consensus       118   ~~~~~i~~~~~s~~l~~lLGF~~~   141 (246)
                         -..+.++ |+. .+|..+||..++
T Consensus       203   LPgCaVD-FT~-SRL~nlLGIrKR   224 (450)
T PF01686.17      203   LPECGVD-FTY-SRINNMLGIRKR   224 (450)
T ss_pred                  eCCcccc-cch-hhHHhhhccccc
```

### Cupiennin

The identity of this gene was assigned based on sequence similarity to other adintovirus Cupiennin genes (which in turn generate hits for cupiennin in HHpred searches, see below)

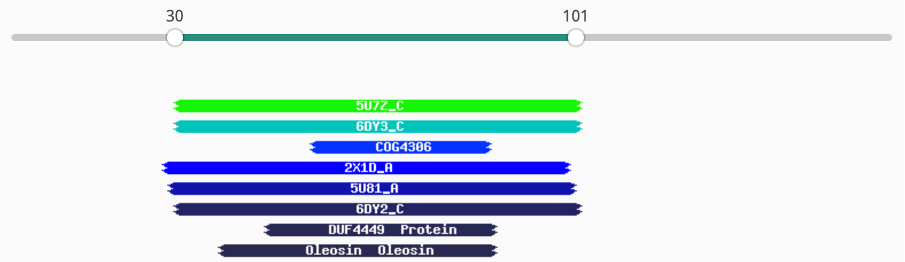


1. **5U7Z_C Acid ceramidase; ceramidase, ceramide, ntn-hydrolase, HYDROLASE; HET: NAG, SO4, BMA; 2.5A {Homo sapiens}; Related PDB entries: 6MHM_C 6MHM_A 5U7Z_A**

Probability: 71.59%, E-value: 15, Score: 28.64, Aligned cols: 70, Identities: 9%, Similarity: 0.016,

```
Q ss_pred                   cccccccccHHHHHHHHHHhHHHHHHHHHHhhh--HHHHHHHHHHHHHHHHHHHHHHHHHHHHccccccCCcccc
Q Q_TerrapeneLip    30    GAPVMYGAGVGGIFRSLFRKAVPLLRRGLEIIK--PHVKTAAQNIAKDVVGHVSRAVLEKVGNTASQEGSGLMY   101 (157)
Q Consensus         30    G~~YQRG~GiGs~l~~LfR~~lPlLksGak~vg--keak~Ag~ni~~Dv~~nLk~kA~~KI~~~~~q~GsG~k~   101 (157)
                        |++++||.-+|..+...++.++.....-....+  ..+.+++...+.+.++.+-....+-|..+  -+|+|+..
T Consensus         45    gsp~erG~q~G~~~k~~I~~~~~~~~~~~~~~~~~~~~~~~~~~~~~~~~~~~~p~~~~Ei~Gi--A~gagl~~   116 (129)
T 5U7Z_C            45    LPPYKRWHELMLDKAPMLKVIVNSLKNMINTFVPSGKVMQVVDEKLPGLLGNFPGPFEEEMKGI--AAVTDIPL   116 (129)
T ss_dssp                 SCHHHHTHHHHHHHHHHHHHHHHHHHHHHHHHSCHHHHHHHHHHHHHHHHCCCCTTHHHHHHHH--HHHHTCCH
T ss_pred                 CCHHHHHHHHHHHHHHHHHHHHHHHHHHHHHHhCCCChHHHHHHHHHHHHHccCCHHHHHHHHHH--HHHhCCCH
```

### Hexon

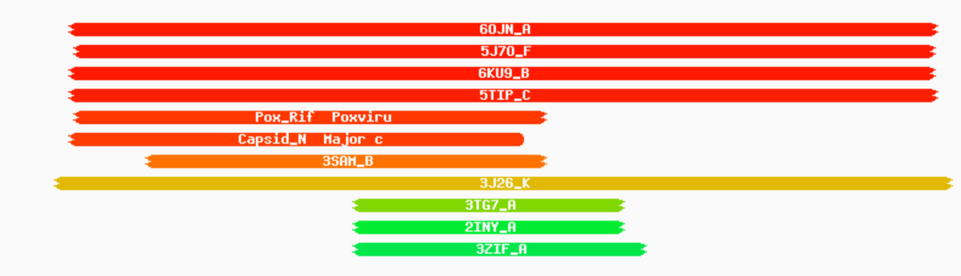


**6OJN_A Major capsid protein; icosahedral, trimers, outer coat, anchor; 8.6A {Singapore grouper iridovirus}; Related PDB entries: 6OJN_C 6OJN_B**

Probability: 97.54%, E-value: 0.0021, Score: 68.79, Aligned cols: 347, Identities: 15%, Similarity: 0.048,

```
Q ss_pred                   ceEEEEECCCCCCCCCCEEEEecCCccceeeccceEEEEEE-EEEcCCCCccccCCceeecccHHHHHHcceEEeCCE
Q Q_TerrapeneHex   30    KSIYIEVPPLSAITESAPIDFFIAGNGIDYMDLNNTLLYLCC-KIVKGDGTELAVDAEVGLVNYPVASIFSQLDVTLGDR   108 (435)
Q Consensus        30    ~~~~~~~~P~s~~~~~~~~i~f~i~~~~~~~y~dl~~s~l~i~~-ki~~~dg~~~~~~~~~v~~~nn~~~slF~~v~~~in~~   108 (435)
                        +.+-++-...+..+-++.+.+.|+..+|   .+.+.+|.|++ .+..............-+++++|.|+++|+++|||+
T Consensus        48    ~~~~~~~~~~~~g~~~fG~~~t~~I~r~GD---Ll~~~~L~i~Lp~i~~~~~~~~~~~~~~~~w~~~lG~~lIe~vel~Igg~   124 (463)
T 6OJN_A           48    KLPTMATRVSGNPAFGQEFSVGVPRSGD---YVLNAWLTLKTPEIKLLETNRLGANGTVRWTKNLMHNAVEHASLTFNDI   124 (463)
T ss_dssp                 BCCCCCCEEESCCCSSCEEEEECCCSSS---BCCCEECEECCCCCCCSCSCCCTTCCCTTSCCCSHHHHCCEEEEEETTE
T ss_pred                 ccceeeEeccCCCCCCCEEEEEcCCCCC---eeeeEEEEEEcCcccccccCCCCCcccchhHHHHHHHHHheeEEEECCE

Q ss_pred                   EeccCCCchHHHHHHHHHhcCChhhhcceeeeeeEEecCCCCCcccccCCCcchHHHheee-------cCCcEEEEEEe
Q Q_TerrapeneHex  109    LISQSNNCYPYRAFIESVLNYSDDTLATQFSAGLFYKDTAGQHEETELDGGNLGFVRRAKLT-------AGSRTVELLGH   181 (435)
Q Consensus       109    ~v~~~~~~~y~y~a~i~~ll~~~~~~~~k~~~l~~~gw~~d~~~~~~~~~~~~~~~~n~G~~~R~~~~-------~~~~~~~~~~~   181 (435)
                        .+++-.   +.-=.+...|..+++.+...-...|...+.....                     .       .......+.+|
T Consensus       125    ~i~~~~~~~~~wl~i~~el~~~~~k~~~~~~~miG~~~~l~~~~~-----------------~~~~~~~~~~~~~~~~l~IP   183 (463)
T 6OJN_A          125    CAQQFN---TAYLDAWTQFNMCEGKRIGYDNMIGNTSDMTNPT-----------------PAQGQDGARTLPSKNLVLP   183 (463)
T ss_dssp                 EEEEEE---HHHHHHTTTTTCCSTHHHHHTTTTSCCCTTCCSS-----------------SSSCCCCCSSSCCCCEECC
T ss_pred                 eeeEec---HHHHHHHHHhhCChhHHHHHHHHcCCCCCCCCCC-----------------CCCCCCcccccCCeEEEEe

Q ss_pred                   ccchhhcCCce-----ecCCceEEEEEEEECCcccceeecCCCCceE------------EEEEEEEEEEEeecChhhhhh
Q Q_TerrapeneHex  182    LHSDLFFQEKL-----LLNGVDVKIKLTRSKDAFCLMGSAAEGFK------------LRIVSASLFVKKVRVAPGVRLG   243 (435)
Q Consensus       182    L~~~~f~q~k-----L~~~~~~~~l~l~rs~d~f~l~~~~~~~~~~~k------------l~I~~~~l~v~~v~~~~~~~~~~   243 (435)
                        |..-|+...+.     -+...+++|++...+-..|+.........            ..+.++.|++..|-++++.+
T Consensus       184    LpFwF~r~~~laLPLiaL~y~ev~i~iklr~~~~l~~~~~~~~~~~~~~~~~~~~~~~~~~~~~~l~~~yv~Ld~eER--   261 (463)
T 6OJN_A          184    LPFFFSRDCGLALPTVVLPYNEIRINIKLRSLQELLVFQNKDTGNVIPISATDIAGGLADTVEAYVYMTVGLVSNVER--   261 (463)
T ss_dssp                 BCCTTSSCSTTCCBSSSCTTCCCEEEEECCSCCTTTSCCSSSCSSCCSSCCTTSSSCSCCCCCBCCEECCCBCCHHHH--
T ss_pred                 ccccceeCCcCccccEEeecCCEEEEEEEeccHHHcEEEecCCCCCcccccccccCccccEEEEEEEEecCCHHHH--

Q ss_pred                   HHHHHhhcceeeeeEEeEEEEEEEe--CCCceeeeeeccccCCCCcEEEEEEecCcccCCCcccCCccCCCCc--------
Q Q_TerrapeneHex  244    HAEALLTANAKYPVDRVGMKVFSI--PAGSRVSNQENLFLGQLPKMLVLGFVDNDAFSGSYAKNPFHFKHYD--------   313 (435)
Q Consensus       244    ~~~~L~k~~~~yp~~~~ei~~~~i~--~~g~~~~~~~~~~~~~~~~~~P~~iiv~~v~~~~~~~g~~~~npf~F~~~~~~~~~~--------   313 (435)
                          +.+.+.+..|-+...+...+..  +.+..+..+.--|.... +.++++++....  .+...+-..+...+
T Consensus       262    --~~fa~~~~~yLI~QvQ~~~~~~~~~~~~~~~~~~~~~~~i~L~f~~PV-k~L~~~~~q~~~~~~--------~~~~y~~~~~~~~~~~~~~~~~~~~   336 (463)
T 6OJN_A          262    --CAMAGTVRDMVVEQMQAAPTHIVNPQNTNNVHVDMRFSHAV-KALFFMVQNVTY--KSVGSNYTCVTPVNGPGNTVME   336 (463)
T ss_dssp                 --HHHTTSCCEEEEEEEEEEEEEECCCCCSSSCCCCCEECCCSSEE-EEEEEEEECSSC--TTCCSCCCSSTTSSSSCSSSCC
T ss_pred                 --HHHhcCchhceEEEEeecCceeeCCCCCCceeeeeeecCce-eEEEEEEeeCcc--cCCCcccceeecCCCCCCCccccC

Q ss_pred                   -------ccEEEEEECCEEccCCCCCCCCCCCChhHHHHHHHHHHhCcccCCCCCcCHHHHccCcEEEEEECCCCccc-c
Q Q_TerrapeneHex  314    -------INFVALYVDGEQIPTKPLQPDFEAGRCVREYMNLVQTAGKHMKDRSLLIDREEFAQGYTLFAFDLSPDQEC-A   385 (435)
Q Consensus       314    -------l~~i~v~~ng~~yP~~~~~~~df~~~~~~~~~~Y~~l~~~~~~~~~~~~~~~~~~i~~~~~f~~~~~~lf~~dls~~~~~~~~~   385 (435)
                               |.++++.+||......            ....|.++.+-...+.....       ..|...+-|-|.|.|.... +
T Consensus       337    ~~~~~~~~i~~~~~l~lNg~~r~~~~---------------~yf~~vqpy~~~~~~p~----------~~giy~YSFaL~p~~~qPs   398 (463)
T 6OJN_A          337    PAMSVDPIKSASLTYENTTRLAN----------MGVEYYSLVQPWYFSASIPV--------YTGYHMYSYALNVGSVHPS   398 (463)
T ss_dssp                 CCGGGSSCCBEEECBTTBCCSCC----------CBHHCCCCCHHHSSSCCCCC--------SSCEEECCCCSCSSCSSCS
T ss_pred                 cccccCCceeEEEEEEECCeecccc----------cChhhHhceeceeeCCCCCC--------CceeEEeeeecCCCCCCCC

Q ss_pred                   ccccceeeeEEEEEEEEecCCCCCC----------------cEEEEEEEEEeCcEEEEc
Q Q_TerrapeneHex  386    DHYSLIKTGNLRAEIRFGKALTV----------------TVNMIVYGVFDNVIEIN   425 (435)
Q Consensus       386    ~~~~~~~~k~g~i~l~i~F~~~~l~~~~~~~~~~~~~~~~~~~~~~~~~~~~~~~~~iv~~~~~d~~~~~id   425 (435)
                        +..+..+=-+.+|++++..+...                 ...+.||++.=+++.|.
T Consensus       399    Gs~NfSri~~~~l~l~~~~~~~~~~~~~~~~~~~~~~~~~~~~~~~~~~~~~~vya~~yNiLri~   454 (463)
T 6OJN_A          399    GSTNYGRLTNASITVTMSPESVVAAAGGGNNNSGYNEPQRFALVVIAVNHNVIRIM   454 (463)
T ss_dssp                 SCEETTSCCCCEEEECCCCCCSCTTSCCCSSCCCTTSTTCCEEEEEEEEEEEEEECC
T ss_pred                 eeEeeccCCcEEEEEEEeCChhceeccCCCCCCCCCcCCCCCEEEEEEEEeceEEEec
```

### Adenain

| | | | | | | |
|---|---|---|---|---|---|---|
| putative thiol protease [Bodo saltans virus] | 60.1 | 60.1 | 55% | 4e-09 | 34.43% | ATZ80653.1 |
| Ulp1 protease [Barrevirus sp.] | 57.0 | 57.0 | 50% | 4e-08 | 32.71% | AYV76948.1 |
| Ulp1 protease [Mimivirus LCMiAC02] | 53.5 | 53.5 | 59% | 7e-07 | 33.05% | QBK89082.1 |
| Ulp1 protease [Indivirus ILV1] | 53.1 | 53.1 | 50% | 9e-07 | 32.08% | ARF09796.1 |
| Ulp1 protease family, C-terminal catalytic domain protein [Mimiviridae sp. ChoanoV1] | 50.1 | 50.1 | 63% | 1e-05 | 27.48% | QDY51658.1 |
| Ulp1 protease [Catovirus CTV1] | 49.7 | 49.7 | 50% | 1e-05 | 35.00% | ARF09252.1 |
| protease [Skunk adenovirus PB1] | 49.3 | 49.3 | 59% | 1e-05 | 30.33% | YP_009162598.1 |
| protease [Bat mastadenovirus G] | 49.3 | 49.3 | 49% | 1e-05 | 30.10% | YP_009325346.1 |

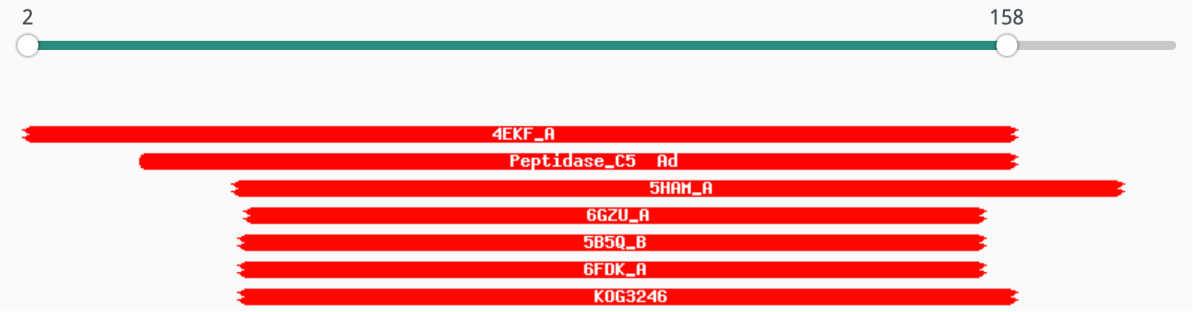


**4EKF_A Adenain (E.C.3.4.22.39); alpha and beta protein (a+b); HET: CSD; 0.98A {Human adenovirus 2} SCOP: d.3.1.7; Related PDB entries: 4WX4_A 1NLN_A 4PIS_A 4PID_A 4PIE_A 5FGY_A 4WX6_C 4WX6_A 4PIQ_A 4WX7_C 4WX7_A 1AVP_A**

Probability: 99.81%, E-value: 1.5e-21, Score: 154.11, Aligned cols: 155, Identities: 19%, Similarity: 0.367,

```
Q ss_pred                 CHHHHHHHHccCcCCCCceEEEEecccCCCCCCCCCCEEEEEECCCCCCCCeEEEEEEEcCCe-EEEEeCCCCCCcc--
Q Q_TerrapeneAde   2      DTVQLSRVLSTDPYTKKNFLDVFPCDWLPGRKLSQRPLGLVVNTHPHNQPGEHWLALYLAERNR-GEFFDSYGHPPNS--   78 (185)
Q Consensus        2      ~~~~i~~~~~~~~~~~~~f~gv~~~~~lp~~~~~~~~~~~~~iN~~~~~~~g~HW~l~~i~~~~~~-i~~fDSlg~~~~~~--   78 (185)
                          |++||++++++.+ ++..|.|||+++ .|......++.+++||+++.+.+|.||++++++...+ ++||||+|..+..
T Consensus        4      s~~eL~~i~k~l~-~~~~FlGvf~~d~~~~~~~~~~~~~~~IvNTd~~~~~G~HWvAl~~d~~~~~~~~yFDSfG~~P~~~~   81 (204)
T 4EKF_A         4      SEQELKAIVKDLG-CGPYFLGTYDKR-FPGFVSPHKLACAIVNTAGRETGGVHWMAFAWNPRSKTCYLFEPFGFSDQRLK   81 (204)
T ss_dssp                CHHHHHHHHHHTT-CGGGEEEEEESS-CCSCCCTTSCEEEEEESCC----CCCEEEEEEETTTTEEEEECTBTTCSHHHH
T ss_pred                cHHHHHHHHHHhC-CCCCeeeeecCC-CCCCCCCCcCeEEEEeCCCCCCCCceEEEEEEeCCCCeEEEECCCCCCcHHHH


Q ss_pred                 ccHHHHHHHHHHH--------cCCCcEEcCCcCCCCCCCCcHHHHHHHHHHHHHC----------------CCCHHHH-HH
Q Q_TerrapeneAde   79     VLFPKSIMKFLNK--------NATDMVFHNRQLQDPRSVACGYHCVFFLHHRSK----------------GLSFDRI-LK   133 (185)
Q Consensus        79     ~~~~~~i~~~L~~~~~~~~~~~~~~~~~~~~~~~~pqQ~ns~dCGvf~l~~~e~~~~~~~~~~~~~~~~~~~~~g~~~~~~~~~~~~   133 (185)
                          +.+......+|++        ...++.++....|+.++..||.||+.|+..+..                |.+...+ ..
T Consensus        82     ~~y~~~y~~~l~r~~l~~~~~~~~~~~~l~~N~~~~Q~~~s~~CG~yCi~FL~~~~~~~~~~~~~~~~~~n~~~~~~~~g~~~~~~~~~~   161 (204)
T 4EKF_A         82     QVYQFEYESLLRRSAIASSPDRCITLEKSTQSVQGPNSAAAGLFCCMFLHAFANWPQTPMDHNPTMNLITGVPNSMLNSP   161 (204)
T ss_dssp                HHHHHHHHHHHHHHCC--------EEEECCSBCSCTTCCCHHHHHHHHHHHHHHCTTSCSSSSTTGGGCCCBCGGGTTCG
T ss_pred                HHHHHHHHHHHHHHHHHHhcCCCcceEEEEcCccccCCCCcchHHHHHHHHHHHHHhCCCCCCCCCcchhhccCCCHHHcCCC


Q ss_pred                 hccccccccHHHHHHHHHHHhhhhhc
Q Q_TerrapeneAde   134    LYSNDLVQNDRMVMNFVKSKFKFLG   158 (185)
Q Consensus        134    ~~~~~~~~~~~~~d~~~~~~~~~~~~~~~~~~~~~~~~   158 (185)
                          .....+..+++++.+|+.++....+
T Consensus        162    ~~~~~~l~~Nq~~ly~~~~~~~~~~~yf~   186 (204)
T 4EKF_A         162    QVQPTLRRNQEQLYSFLERHSPYFR   186 (204)
T ss_dssp                GGHHHHHHHHHHHHHHHHHHHCHHHH
T ss_pred                CChhhHHHHHHHHHHHHHHHHhCHHHH
```

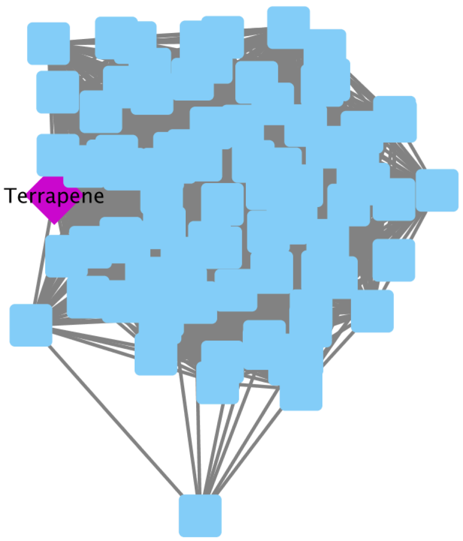


### Adintovirus conserved protein category 2 (Adintoc2)

| Description | Sequence Length | Sequence |
|---|---|---|
| [illegible] | [illegible] | [illegible] |
| [Adintoc2@Sinocycloheilus_Adinto_LAVE01068915_4_2312_-_2716_] | 135 | LRTDLDQREKAK... |
| [Adintoc2@Sinocycloheilus_Adinto_LAVE01068915_5_2720_-_2959_] | 80 | SEFLQSFAELNIPF... |
| [Adintoc2@Stegodyphus_spider_roughly_circular_22kb_Adinto_retrovirus_AZAQ01114673_32_8111_-_7776... | 112 | VYSYLLLFLSLVLL... |
| [Adintoc2@Strongylocentrotus_urchin_intact_Adinto_AAGJ05080195_11_9821_-_10564] | 248 | IRMENTRKMVLVP... |
| [Adintoc2@Symphodus_wrasse_Figure_intact_Adinto_(nearby_ORFB_RT_is_host_LINE)_MWVA01005046_31_2... | 320 | TKIIFLFLKKSSEM... |
| [Adintoc2@Symphodus_wrasse_intact_Adinto_ITR_Hexon_Integrase_in_Blast_MWVA01004576_30_22484_-_2... | 322 | TKIIFLFLKKSSEM... |
| [Adintoc2@Terrapene_BK010890] | 239 | MEYAEKMFLVPS... |
| [Adintoc2@Torymus_wasp_9kb_Adinto_frag_UCWB01007878_8_6364_-_5549_(REVERSE_SENSE)_] | 272 | NNGESKKMVLISK... |
| [Adintoc2@almost_intact_Adinto_AOMJ02232319_5_2433_-_3209] | 259 | LLNDLFIVTFVTS... |
| [Adintoc2@intact_Adinto_ITR_direct_repeats_AYNC02010475_9_12999_-_13736] | 246 | DRFWVLLHYRTM... |
| [Adintoc2@intact_Adinto_no_ITR_AZAQ01040193_15_3257_-_2778_(REVERSE_SENSE)] | 160 | MEHAKKYVLMPE... |

### GasderminX

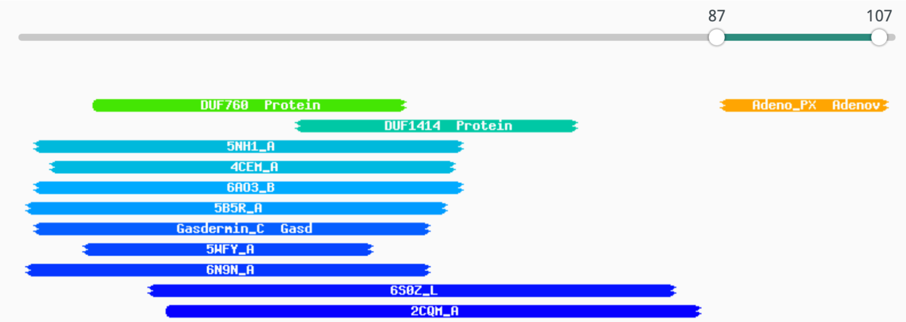


**5NH1_A Gasdermin-D; apoptosis, Gasdermin, pyroptosis, inflammasome, innate; 2.04A {Homo sapiens}; Related I 5WQT_B**

Probability: 57%, E-value: 53, Score: 26.69, Aligned cols: 53, Identities: 15%, Similarity: 0.107,

```
Q ss_pred                 hHHHHHHHHHHHHHhcCCHHHHHHHHHhCCHHHHHHHHHHHHHHHHcCCCCCCHH
Q Q_TerrapeneGas   3     SCVKRNLGLLKLLSKSSPQQRRAILCSASDDLVAAISEIALNTLKGNVPLTQN    55 (109)
Q Consensus       3     ~~vk~~~~~L~~L~~~~~~~qRkaLLk~a~~~~~l~~l~Ec~lNiLkG~v~Ls~~    55 (109)
                        +...--+-++.+|..+++.|..-|-..+..+.+....|.+.+|++.+++.+++
T Consensus       89    ~l~~ai~~ll~AL~eLse~q~~LL~~~~e~~iL~~~~~LV~~ile~~~~~~~~~    141 (207)
T 5NH1_A          89    ELAIPVVYLLGALTMLSETQHKLLAEALESQTLLGPLELVGSLLEQSAPWQER    141 (207)
T ss_dssp                 HHHHHHHHHHHHHTTSCHHHHHHHHHHHHTTCCHHHHHHHHHHHHHHTCSTTSC
T ss_pred                 HHHHHHHHHHHHHHhCCHHHHHHHHHHHhcCcchHHHHHHHHHHHhcCccccc
```

**PF05829.12 ; Adeno_PX ; Adenovirus late L2 mu core protein (Protein X)**

Probability: 85.76%, E-value: 1.6, Score: 27.65, Aligned cols: 17, Identities:

```
Q ss_pred                 cCcccHHHHHHHHHHHHHHhh
Q Q_TerrapeneGas   87    SGGFIGPLLSFAIPLITGLLT    107 (109)
Q Consensus       87    ~gGfl~~ll~~l~p~l~~ll~    107 (109)
                        +|||    +++++|++.+.+|
T Consensus       11    rGGf----lpaLiPiiaaaIg    27 (42)
T PF05829.12      11    RGGF----LPLLVPIIAAAIS    27 (42)
T ss_pred                 cccc----HHHHHHHHHHHHHH
```

Note: this result is for the single GasderminX sequence of the example virus. Multiple alignments of related GasderminX genes give higher probability hits in HHPred searches

### FtsK

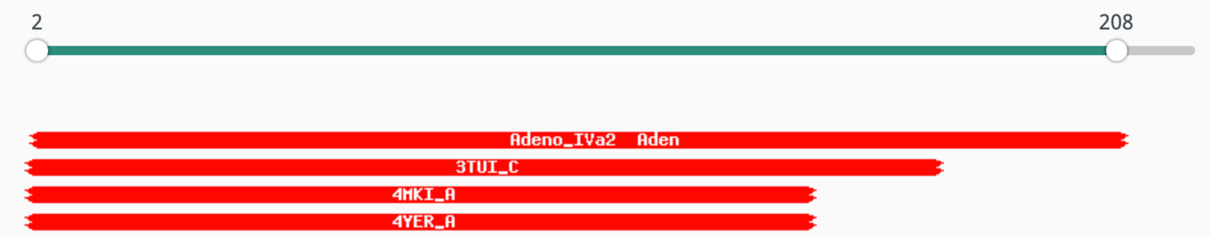


**PF02456.15 ; Adeno_IVa2 ; Adenovirus IVa2 protein**

Probability: 99.44%, E-value: 2.3e-14, Score: 117.25, Aligned cols: 202, Identities: 15%, Similarity: 0.164,

```
Q ss_pred                 CccccCCeEEE-EECCCCCCHHHHHHHH--HHhhhcccccCcccEEEEcCCchHHHHHHHHHCCCcEEEcCCCCCCCccc
Q Q_TerrapeneFts  2     DTRWKHPFSAI-LAGPSNCGKSYFIKNV--LDNAKHTLSVMPENIVWCYSCWQPLYKELLCKYPFINFVEGLPDAFDDDS    78 (223)
Q Consensus       2     ~~~i~~g~~~~-i~G~~GsGKStll~~l--------------g~i~~~~~~~~~~~~~~~~~~~~~~~i~~~~~~~~~~~~    78 (223)
                        ||++..|+.++ |+|+||||||||+++|  ++    ...+..|.|.+.+..........-.......-...... .....
T Consensus       73    sl~i~~g~~i~~I~G~nGsGKStLL~~L~~~g-------p~~G~I~~~g~~~~~~~~~~~~~~~~~~~~~~~~~~~~~~~~    147 (364)
T PF02456.15      73    SLNYGLNPFIGIISGPTGTGKSQLIRNLISNK----LISPPPETVIFITPTKGMLSPEETNLWKLQLQEGNYEK-NTDGI    147 (364)
T ss_pred                 ceEEecCCEEEEEECCCCCCHHHHHHHHHHCC----CCCCCCCeEEEcCCCcccCCHHHHHHHHHhhhcCCCcc-CCCCC

Q ss_pred                 cCCC-----------------------------------CCceEEEEecccCCcc--CCHHHH--HHHHHHhcc--CCeE
Q Q_TerrapeneFts  79    LFPT-----------------------------------NKVNMIIIDDLMNSAC--ESDEIE--KAFTKYVHH--RNLS    117 (223)
Q Consensus       79    ~~~~-----------------------------------~~~~llilDd~~~~l~--~~~~~~--~~~~~~~~~--~~~t    117 (223)
                        +...                                   .+|.++|+|||+.++|  ....+.  .++..+.+.  .+.|
T Consensus       148   v~~~~~~~~~~~~~~~~~~~~~~~l~~~~qr~~la~al~~~p~llllDEp~~~LD~~~~~~~~~~~~ll~~l~~~~~~g~t    227 (364)
T PF02456.15      148   CPITSVFSVDFVEIDFETAVSPDNLDINNDKSCFVQAAKNGNVCIVIDECMKCLIDKRNISPLFCSLPSKISSKFKTGFS    227 (364)
T ss_pred                 CcccccccchhhcCCCHHHccChhhcCCCcHHHHHHHHHHcCCEEEEEEcchhcCCchhchHHHHhhhhHHHHHHhcCCcE

Q ss_pred                 EEEEEehhhhcCC-CcceeHHh-ccEEEEecCcccHHHHHHHHHHHCCC-CHHH---HHHHHHHHHhCCCCEEEEECCCC
Q Q_TerrapeneFts  118   IMYIVQNVFCQGK-KSRTINLN-TKYMVLFKNPRDKLQIATLARQMYPG-KAQF---FLEAFEDATKRPYGYLVVDLNAS    191 (223)
Q Consensus       118   ~i~~s~~~~~~~~-~~~~i~~~-~~~v~~~~~~~~~~~~~~~~~~~~~~-~~~~---~~~~~~~~~~~~~~~~~i~~~~~    191 (223)
                        +++++|+...... +++.+... ...+...+......+....+..+... +...   +.+.+......+|++++.+..+.
T Consensus       228   vii~th~~~~~~~~~~d~i~~L~~~g~i~~~g~~~~~~~~~~~~~~~~~~~~~p~~~~~~ll~~i~~~~~~~~~~~~~~y~~~p~    307 (364)
T PF02456.15      228   MFVVLHNINPTSGNGNNIMDLKIQAKLHILSSKNAPFQLSRFVNTYGTGMSPQLKAVLMNSIIEESKNDYSFILFNVEPP    307 (364)
T ss_pred                 EEEEeCChhHhcccCCEEEEEeeCCEEEEccCCCcccccchHHHHHhccCCCHHHHHHHHHHHHHhhhcCCCceEEEecCCCC

Q ss_pred                 CCccccc-ccCCCCCCCC
Q Q_TerrapeneFts  192   TPEAYRL-RTGLFPPDWP    208 (223)
Q Consensus       192   ~~~~~~~-~~~~~~~~~~    208 (223)
                        .....+. ......+..|
T Consensus       308   ~~~~~w~~~~~~~~~~~p    325 (364)
T PF02456.15      308   RPGFSWCALLNGGNCCEP    325 (364)
T ss_pred                 CcccccchhhcCCCCCccc
```

### PolB

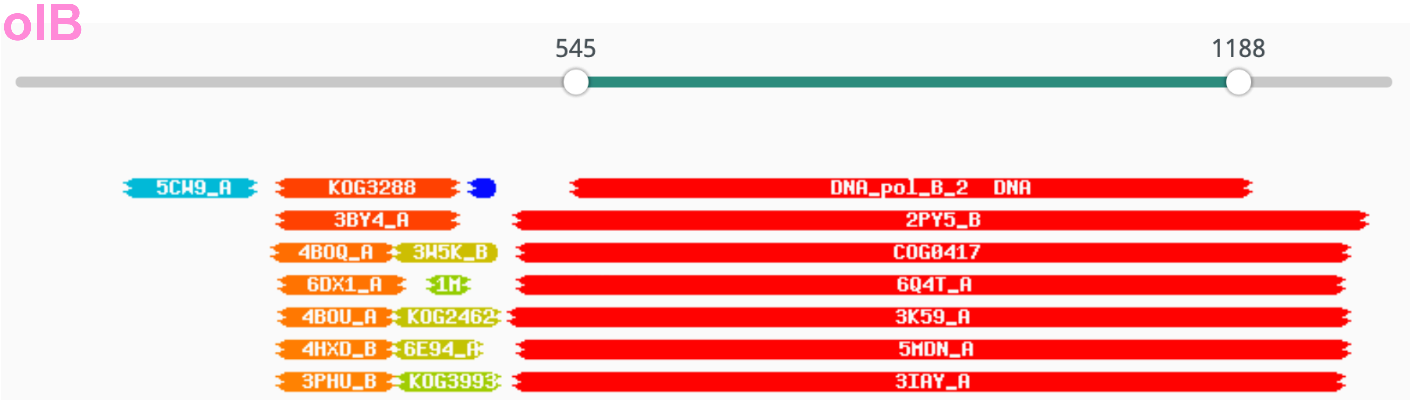


Template alignment | CDD

1. **PF03175.13 ; DNA_pol_B_2 ; DNA polymerase type B, organellar and viral**

Probability: 100%, E-value: 2e-47, Score: 470.57, Aligned cols: 426, Identities: 23%, Similarity: 0.332,

```
Q ss_pred                   cccCcEEEecCCCCccHHhHHHHHHH-cC---------CCceEEecCCeEEEEEEE------eCcEEEEehHhccCcchH
Q Q_TerrapenePol   545     EFRDYTFLAHNSKGYDAYFIVRQLLK-EK---------MGLELITQGSKLMCVEVK------ALGIRFIDSLNFLPMKLS   608 (1337)
Q Consensus        545     ~~~~~~~viaHN~~~YD~~fIl~~l~~~~~~~~~~~~~~~~~~~~~~~ii~~g~kii~~~~~~~~~~~~~~~~~i~F~DS~~fl~~sL~   608 (1337)
                            ++++.+|+|||+++||..||++.+.+ .+          ..+.++.++++++.+++.      ..++.|+||++++|.||+
T Consensus        6       ~~~~~~i~~hN~~~fD~~~i~~~~~~~~~~~~~~~~~~~~~~~~~~~~~~~~~~~~~~~~~~~~~~~i~f~Ds~~~~~~sL~   85 (457)
T PF03175.13       6       KYRNVLFYAHNLGKFDAVFLLKELLRFNLTTEGLNNPYEFPAPLTRNSDILKLTIKRTVDGKTRKVTIMDSFALLPRSLR   85 (457)
T ss_pred                   ccCCeEEEEEcCCCcCHHHHHHHHHHccCCccCCCCCCCCCceeeeCCEEEEEEEEEcCCCCeeEEEEEehhHhccccHH


Q ss_pred                   HhHHHhCCCCccccCCccccCccccCCCCCCCChHhcCCcCCCHHHHHHHHHHHHHcCCCCCCHHHHHHHHHHHHHHHHH
Q Q_TerrapenePol   609     KLPQAMGFEGCKGYFPHFFNTLENQNYVGPMPGMEHYGVESMMPREKAEFLDWYRDHSSEAFDLQKELAYYCQQDVKILR   688 (1337)
Q Consensus        609     ~l~k~fgl~~~Kg~FP~~fn~~~n~~~~~~~lP~~~~Y~~~~m~~~~~~~~f~~wy~~~~~~~F~~~~el~~Yc~~DV~iL~   688 (1337)
                            +++++||++..|+.|||.++++.+..+.+++|+..+|.++.+  ++.++|.+|        .|++.+++..||.+||.+|+
T Consensus        86      ~~~~~~~~~~~K~~~p~~~~~~~~~~~~~~~p~~~~~~~~~~~~~~~~~~~~~~~~~~~~~~~~~~~Y~~~Dv~ll~   156 (457)
T PF03175.13       86      DLCKDYNIDNPKSYFPYLFSVDSHLFYTGATPNINYYLDIPK--AEYLKLVKD-------DWDFKDETLEYLNLDLISLH   156 (457)
T ss_pred                   HHHHHcCCCCCcccCChHHccccccccCCCCCCccccCCCCH--HHHHHHhhc-------CCCcHHHHHHHHHHHHHHHH


Q ss_pred                   HHHHHHHHHHHHHhccCCccccCCCCCccccccCcChhhhhHHHHHHHHHHHHHhccCCCcc--cccCCCCcccccccCHH
Q Q_TerrapenePol   689     QACILYRKEIMKMTEKGDVVEKEPGKFTEIKLCIDPFRYITLASVCMAMYRFMFLEPNTV--ALLPPDNYHRQKKRYSTP   766 (1337)
Q Consensus        689     ~~~~~fr~~~~~~~~~~~~~~~~~~~~~~~~~~~~~~dp~~~~Tias~~~~i~r~~~l~~~~i---~ip~~gy~~~~~~~S~~   766 (1337)
                            +++.+|++.+.+.+                     ++||+...|+||+++.++++.++.+...  ..+|
T Consensus        157     ~~~~~~~~~~~~~~~~~~~~~~~~~~~~~~~~~~~~~~~~~t~~s~a~~~~~~~~~~~~~~~~~~~~~-------------   204 (457)
T PF03175.13       157     QVLVKVNKAINFLF-----------------DIEFTSCATVSSLANKIFLSKFYDDKNKAIPLVK-------------   204 (457)
T ss_pred                   HHHHHHHHHHHHHh-----------------CCCcccccHHHHHHHHHHHhccCCCCccCCccC-------------


Q ss_pred                   HHHHHHHHHhccCCceEecccCCceEecceEEeceeccCCceEEEEEeCceecCCCcccCCCCCCCCCCCCCHHHHHHHHH
Q Q_TerrapenePol   767     SIQWLLYISHKENIQIRHALQGGELQVGPYFLDGYANVDGVRTAFEFNGCFFHGCVTCYCEKAQNPMTGTSFGFLYYKTQ   846 (1337)
Q Consensus        767     al~wL~~~~~~~~~~~i~h~~~ggE~~i~~~~vDGy~~~~~~~~v~E~~GC~~HgC~~C~~~~~~~~~~~t~~~l~~~t~   846 (1337)
                                                 +.++..++            |++++|||++++...+.....+.|.++|+|||||
T Consensus        205     --------------------------------------------------------------------------------   204 (457)
T PF03175.13       205     --------------------------------------------------------------------------------   204 (457)
T ss_pred                   --------------------------------------------------------------------------------


Q ss_pred                   HHHHHHHHcCCEEEEeehhhHHHHHHhcCHHHHHHHHhCCCCCCCCcchhhhcccccceeeeeCCCCCCeEEEEeCccccH
Q Q_TerrapenePol   847     LKTDYLKRLGFVVRTLWEHEWEVMKETDRELADFLNRAQLPQPLVPRDALFGGRTNAIRLYYKPNPGEEIHYYDFTSLYP   926 (1337)
Q Consensus        847     ~r~~~l~~~g~~v~~~Wece~~~~l~~d~em~~f~~~~~~~~~~l~~R~a~~GGRt~~~~~~~~~~~~~~i~y~DvnSLYP   926 (1337)
                            ++|..  .+|.|.+.+..........  .|++.|++.+|.+..+|++|.+.+++|+||++.
T Consensus        205     ---------------------------~~~~~~~~~~~~~~~~~~~~~~~~~GG~~~~~~~~~~g~~~~~i~~~D~nSlYP   246 (457)
T PF03175.13       205     ---------------------------DIDLFNDI-----------HEAYYGGRVEVFNPIIMADSTKSYYYYDVNSLYP   246 (457)
T ss_pred                   ---------------------------CccHHHHH-----------HHhhcCccceeceeeeecCCCCeEEEEechhhCH


Q ss_pred                   HHHhcCCCCCCCCceeecCCCCcccC--eeEEEEEEeCCcccCCCCCCceeCCeEEecCChhccHhcCccccCCChhhhc
Q Q_TerrapenePol   927     FVNKTKEYPIGHPDIVYDKFGPLANY--FGIAKVKVYPPRGLFFPLLPVRVGGKLMFPLCRTCAETEQRETCTHTDEERA   1004 (1337)
Q Consensus        927     ~vm~~~~~P~G~p~~~~~~~~~~~~~~~~--gi~~v~v~~P~~l~~PvlP~r~~~kl~fpl~~~c~~~~~~~~~~c~h~~~~r~   1004 (1337)
                            +.|+++++||+.|++.|++ .++++++.|++. .++|++||+.++++|.+
T Consensus        247     s~m~~~~p~~~~~~~~~~~~~~~~~~~~~g~~~~~~~~~p~~~~~p~~p~~~~~~l~~~~~~-------------------   306 (457)
T PF03175.13       247     FASIN-DIPGLKCTFYEVVKANVNIHELFGFFYCKIKSPDNLYLGLLPKRTETSLIFPGGA-------------------   306 (457)
T ss_pred                   HHhcc-CCCCCCcEEeecccccccccccceEEEEEECCchhcccCCccccCCceeCCCcc-------------------


Q ss_pred                   eEEEEehHHHHHHHHCCCEEEEEeEEEEechhcChhHHHHHHHHHHHHHHHhcCCCccccCCHHHHHHHHHHHHHHhCCCCC
Q Q_TerrapenePol   1005    ILGTWCTVELNAAIAKGYAVAKIYEIWHFNEKSDKLFSEYIKLHLRQKQEASGYPSWCTDEEKQNKYVSDFYQKEGVHLR   1084 (1337)
Q Consensus        1005    ~~g~~~s~ELk~Al~~Gy~i~kI~e~~~f~~~~~~lF~~yI~~~~~~K~easg~p~~~~~~e~~~~yi~~~~~~gv~ld   1084 (1337)
                                 .+..+++.++||+|||+||||||++..+++.++++.+.     ........+.....+++....+.+........  .
T Consensus        307     ~~~~~~~~el~~a~~~g~~~v~~~~~~~f~~~~~~~~f~~~i~~~~~~R~~------------------------------   354 (457)
T PF03175.13       307     WEGWYFSEELKFAVAHGYE-VEIIKGYNFDRV-SNVFNEFVQEVYKVKCN-----------------------------   354 (457)
T ss_pred                   eeceecHHHHHHHHHCCCE-EEEEEEEEecCC-CchHHHHHHHHHHhhC-----------------------------


Q ss_pred                   HHHhcCCHHHHHHHHHHHHHHHHHHhccCCCCCCeEEeCCHHHHHHHHhCCCceeeEEEEecCCeEEEEEEeccccccC--C
Q Q_TerrapenePol   1085    QHEIRSNPAKRQIAKLFLNSLWGKFGQRSNLPNTSIVRDPDELLQYLFSPNYEVSSCEFIDDETACVSWKHAKERYS--V   1162 (1337)
Q Consensus        1085    ~~k~~~N~~~r~i~KL~lNSlyGKf~q~~~~~~~~~~~i~~~~~~~~~~~~l~~~~~~~~~~~~~~~~~~~~~v~~~~~~~~~~~~~~---   1162 (1337)
                            ..+++++||++||++||+.|+++|..
T Consensus        355     -----k~~~~~~~~Kl~~Ns~yGk~~q~~~~~~~~~~~~~~~~~~~~~~~~~~~~~~~~~~~~~~~~~~~~~~~~~~~~   427 (457)
T PF03175.13       355     ----PNNVTEKNIAKLILNSLIGRFGMNINKIKTSLVPSEKH---NELLTTRVVKSTTDLGSGMFMDIYVPGIDKNICED   427 (457)
T ss_pred                   ----CCCHHHHHHHHHHHHHHHHHHHhhcCCCCCeEEEeChHHH---hHhccCCeeeeEEEcCCCeEEEEeecCCCcccccC


Q ss_pred                   CCCCCHHHHHHHHHHHHHHHHHHhh
Q Q_TerrapenePol   1163    SGNTNVFIACFTTAYARLELYSLLDG   1188 (1337)
Q Consensus        1163    ~~~~ni~ia~~iTs~ARl~L~~~m~~   1188 (1337)
T Consensus        428     ~~~~~i~i~~~i~~~ar~~~~~~~~~   453 (457)
T PF03175.13       428     FNLDYIEVLNSTNTDEKGTMPDNNVS   453 (457)
T ss_pred                   CCCccHHHHHHHHHHHHhhhhHHHHh
```

### PolB (continued)

**KOG3288 OTU-like cysteine protease [Signal transduction mechanisms, Posttranslational modification, protein turnover, chaperones]**

Probability: 94.53%, E-value: 0.2, Score: 61.1, Aligned cols: 165, Identities: 9%, Similarity: -0.006,

```
Q ss_pred                ceeEecccCCCCcHHHHHHHHHHccCCCCc-HHHHHHHHHHHHH------------------hCCC-cCCCCCHHHHHHH
Q Q_TerrapenePol   263   QCLVDLTYTGTNLCFAGGLLAVMSDRKPTD-AELLAEARKLHEK------------------LGWS-DQKKVMLSDVAKF   322 (1337)
Q Consensus       263   ~~lI~i~n~~Dn~Cf~~AI~~~l~~~~~~~~~~~~~~~a~~l~~~~~~~~~~~~~~~~~~~~~~gi~~~~~~~~l~Di~kf   322 (1337)
                         ..++.....+|+.||.+||..++....... ..+.+.....+..                  ..|. ...-.+--+|..+
T Consensus       108   ~~~~~~~v~gDGnCLFrAis~~l~~~~~~~~~~~LR~~v~~~i~~n~~~y~~~i~g~~~~~Y~~~~~~~~~WGg~iEl~al   187 (307)
T KOG3288         108   GVLSRRVVPDDNSCLFTAIAYVIFKQVSNRPYELREIIAQEVASNPDKYNDAILGKPNKEYCAWILKMDSWGGAIELSIL   187 (307)
T ss_pred                ceEEEEEeCCCCCcHHHHHHHHHhcCcCCCHHHHHHHHHHHHHHCHHHhHHHhcCCCHHHHHHHHhcCCccCCHHHHHHH

Q ss_pred                HHHHCCCeEEEEEECCCCceEEecCCCCCCcEEEEEEeCCCchhcccHHHHhCCCccccccCCccCC--CCccccccccc
Q Q_TerrapenePol   323   EQHLGVNIQVVLYTAKGGWGFFKTGGTVYPKTYFILLHDEHYYGVLDVKKLFGAKNYCEFCHMVYSH--DHSCRYHCRLC   400 (1337)
Q Consensus       323   e~~l~~~~~I~Vy~~~~~~~~~~~~~~~~~~~~~~~~i~Lll~~~HY~~I~~~~~l~~~~~yC~~C~~~y~~--~H~C~~~C~~C   400 (1337)
                         .+.+  +..|.||.........+.........|.|+|..+||+.+.....  ........+.-..+.  -......|...
T Consensus       188   a~~~--~~~I~v~~~~~~~~~~~~~~~~~~~~~~~~i~L~y~g~HYd~l~~~~~--~~~~~~~~~f~~~d~~~~~~a~~l~~~~   263 (307)
T KOG3288         188   SDYY--GVEICVVDIQTVRIDRFGEDKNFDNRVLLLYDGIHYDPLAMNEF--KPTDVDNTIFPVSDDTVLTQALQLASEL   263 (307)
T ss_pred                HHHH--CCeEEEEECCCCeEEEeCCCCCCCCeEEEEEcCccceeeeecCC--CCCCCCceeccCCcHHHHHHHHHHHHHH

Q ss_pred                CCc-cCCCCCCCeEECcccCcEeCCHHHHHHH
Q Q_TerrapenePol   401   LSV-TCSDSVGVQQRCPSCRLYCRSKECLDRH   431 (1337)
Q Consensus       401   ~~~-~C~~~~~~~~~~~~~C~~C~~~F~sq~Cfd~H   431 (1337)
                         ... .-+......++|..|+..+.++.--..|
T Consensus       264   ~~~~~~td~~~f~l~C~~C~~~~~~g~~~a~~h   295 (307)
T KOG3288         264   KRTRYYTDTAKFTLRCMVCQMGLVGQKEAAEH   295 (307)
T ss_pred                HhcCCcccCccceEEecccCCeecCHHHHHHH
```

### Oncoid

The ORF designated “oncoid” is named based on synteny with other adintovirus cysteine-rich ORFs that encode short linear motifs of known oncogenes. The oncoid ORF in this example does not itself show clear hits for the linear motifs detailed in the text.

### Reference-guided re-assembly and identification of termini

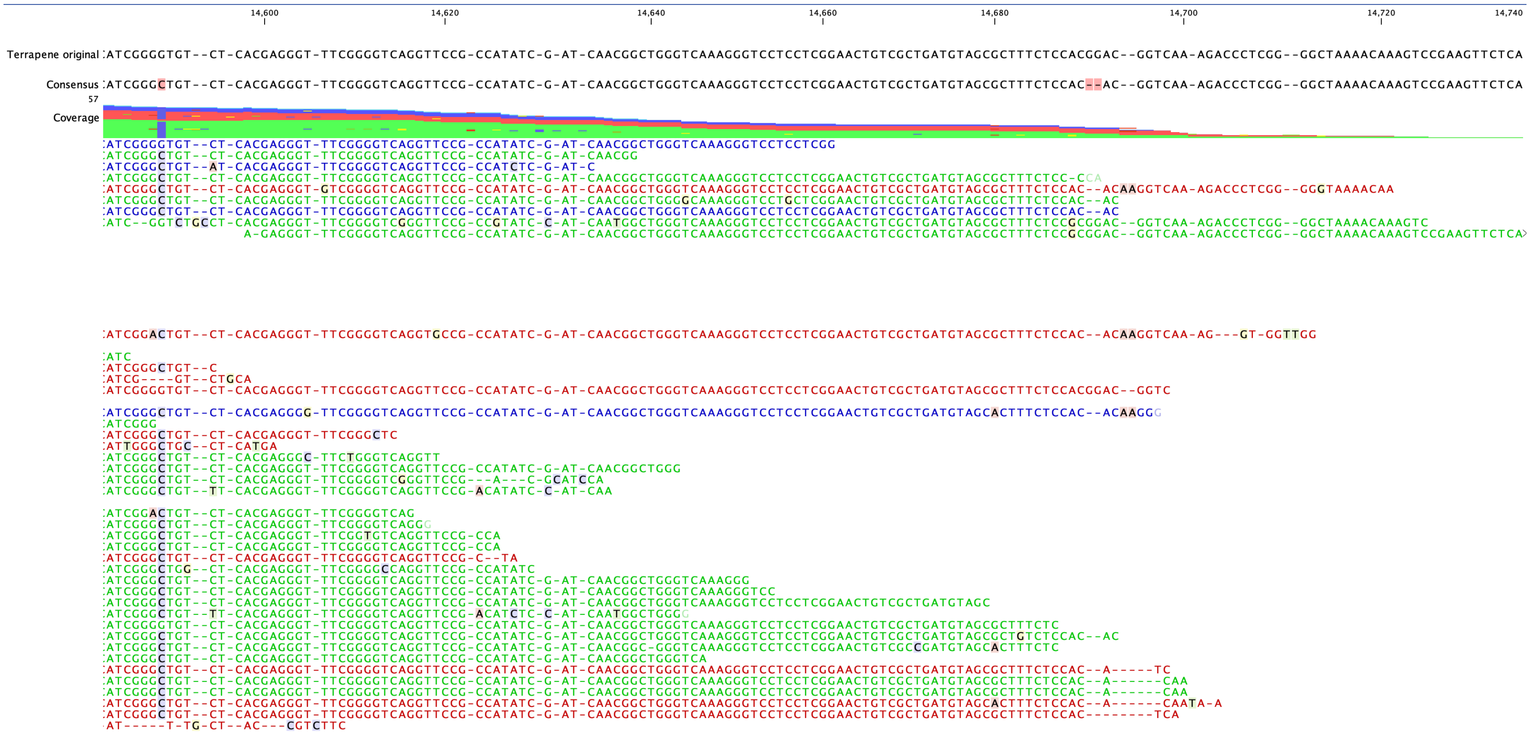

### Examples of other genes

#### Mayetiola PLA2X

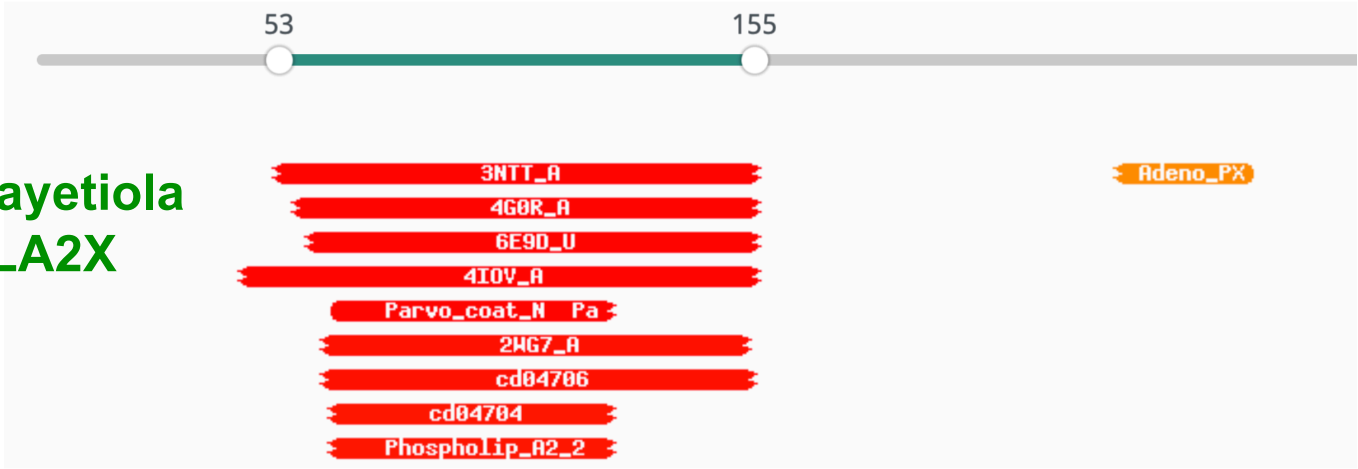


**1. 3NTT_A Capsid protein; Adeno-associated virus 5, gene therapy; 3.45A {Adeno-associated virus - 5}**

Probability: 99.47%, E-value: 1.1e-16, Score: 170.57, Aligned cols: 89, Identities: 25%, Similarity: 0.321,

```
Q ss_pred                 HHHHHhCCccccCCCCCCCCCCCchhHhHhcCCCCCCHHHHHHHHHhhhhccCCCC-----chhHHHHHHHHHHHHHHHHhc
Q Q_MayetiolaPLA    53   DKIIDKLPFELHVPKYQYCGPGTKLDKRLARGDPGINPLDVACKQHDIAYTEHPH-----SDDRYVADKTLQKAAMKRVF   127 (315)
Q Consensus        53   n~~in~lp~elhlPGYnyCGPGt~l~kRL~rGd~pIN~LD~aCkeHDicY~~~~~~~~~~~~~~~R~~aDkiL~~~A~~rv~   127 (315)
                         |.+++..+.++|||||||||||.++     .+.|||.||+||++||+||++..+     +..+|.||++|+++++..
T Consensus        34   ~~~~~~~~~gl~lPGYnYcGPGn~l~-----g~Pin~~D~acr~HD~~Y~~~~~~g~npy~~~~~aD~~fi~~l~~~~--   106 (724)
T 3NTT_A           34   NQQHQDQARGLVLPGYNYLGPGNGLD-----RGEPVNRADEVAREHDISYNEQLEAGDNPYLKYNHADAEFQEKLADD--   106 (724)
T ss_dssp               --------------------------------------------------------------------------------
T ss_pred               ChhhcccCCceecCCCCcCCCCCCCCC-----CCCCCCHHHHHHHHcchhHHhHhcCCCccccCChHHHHHHHHHHcC--
```

```
Q ss_pred                 CCCCChhHHHHHHHHHHHHHHHHHhccC
Q Q_MayetiolaPLA   128   SKDASFGERATALGVAAAMKAKRTLTSR   155 (315)
Q Consensus       128   a~dsslgEKaaA~aV~~aMK~K~klGmg   155 (315)
                                ..+.++++..+|++|++++++
T Consensus       107   -------~s~~g~~~~~~f~~K~~i~~~   127 (724)
T 3NTT_A          107   -------TSFGGNLGKAVFQAKKRVLEP   127 (724)
T ss_dssp               ----------------------------
T ss_pred               -------CChHHHHHHHHHHHHHHHhcc
```

**51. PF05829.12 ; Adeno_PX ; Adenovirus late L2 mu core protein (Protein X)**

Probability: 87.93%, E-value: 0.4, Score: 35.25, Aligned cols: 30, Identities: 43%, Similarity: 0.656,

```
Q ss_pred                 cCccccc-hh-hhcchhHhhHHHHhhHHHHHHHH
Q Q_MayetiolaPLA   231   SGGVLPL-IP-IFAGLSALGTIAGTTTSVISA   260 (315)
Q Consensus       231   ~Gg~Lp~~-ip-ifagLsA~galaGGaa~iakA   260 (315)
                         +|||||+ +| |-++++|+...||-+-.+.++
T Consensus        11   rGGflpaLiPiiaaaIgAiPgIAs~ai~a~n~   42 (42)
T PF05829.12       11   RGGFLPLLVPIIAAAISAAPGIAGAVLAAKNH   42 (42)
T ss_pred               ccccHHHHHHHHHHHHHHhhhHHHHHHHHHHhcC
```

#### Astyanax Cupiennin

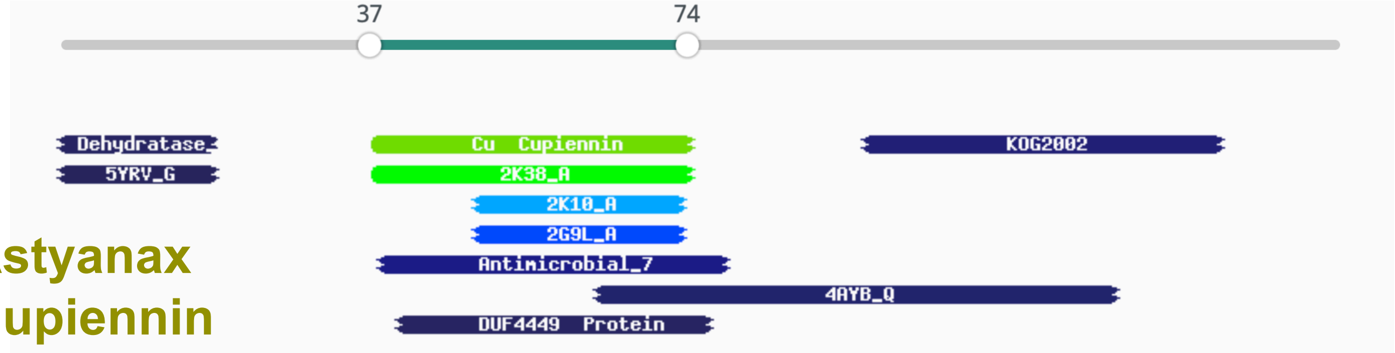


**1. PF17563.2 ; Cu ; Cupiennin**

Probability: 76.67%, E-value: 18, Score: 23.9, Aligned cols: 31, Identities: 23%, Similarity: 0.43,

```
Q ss_pred                 chhHHHHHHHhhhHHHHHHhhHhhHHHHHHHHHHHHHHHH
Q Q_AstyanaxCup     37   GFGGIFKALFRMAVPLIKRGITIVKPHLKTAAKGIVTD   74 (150)
Q Consensus         37   GiGsil~gLfR~~lPl~~rgakaVGKEAl~aG~~Il~D   74 (150)
                         |+|++|.-|..       ..|+.|.++|...|..-+..
T Consensus          1   gfg~lfkflak-------kvaktvakqaakqgak~va~   31 (35)
T PF17563.2         1   GFGSLFKFLAK-------KVAKTVAKQAAKQGAKYIAN   31 (35)
T ss_pred               CchHHHHHHHH-------HHHHHHHHHHHHHhHHHHHh
```

Branchiostoma Bcl2

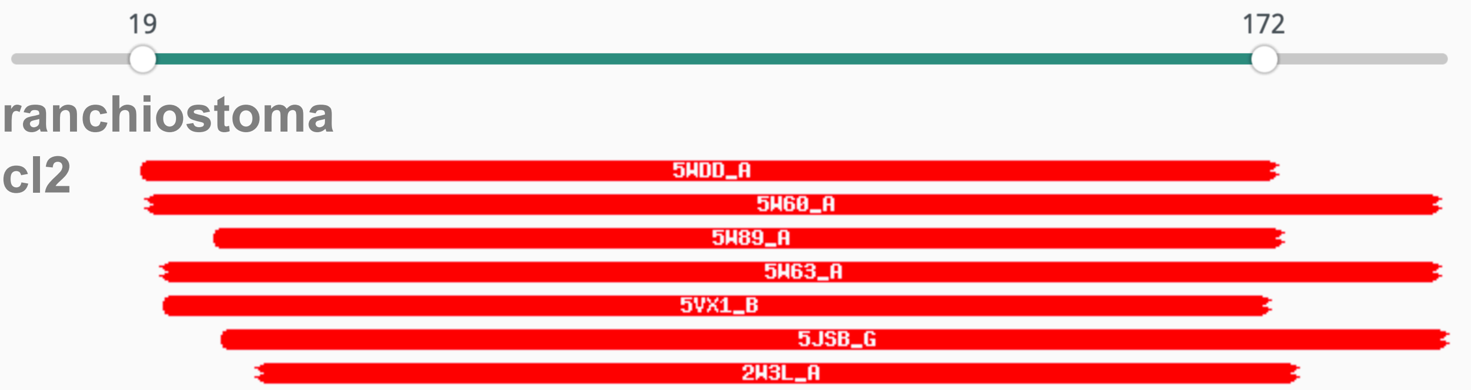


7. **2W3L_A APOPTOSIS REGULATOR BCL-2; APOPTOSIS; HET: DRO; 2.1A {HOMO SAPIENS} SCOP: f.1.4.1; Related PDB entries: 2O2F_A 2W3L_B**
Probability: 99.96%, E-value: 6.7e-31, Score: 204.21, Aligned cols: 139, Identities: 16%, Similarity: 0.266,

```
Q ss_pred                HHHHHHHHHHHHHhcccCCCC-CCchHHHHHHHHHHHHHHHHHcchhHHHHHHHHHhcCC-CChHHHHHHHHHHHHHhcC
Q Q_BranchBcl2    35     QTEDIFRHLLHHRAPNVFPRN-ERRENRAIYRKVVQVSNDVTERKEWADLLCCFVSTTTT-TNGASRLRALLENVLNEDG   112 (197)
Q Consensus       35     qt~~l~~~yi~~~l~~~~~~~-p~~~~~~i~~~Lr~lgdele~~~~~~~~f~~~~~~l~~-~~~~~~~f~~v~~elf~dg   112 (197)
                         +++.|+.+||++++.+++... +++++.+++++|+++|++++++|  +..|++++++++. +..++..|.++++++|.||
T Consensus       2      ~~~~l~~~yi~~~l~~~g~~~~~~~~~~~~~~~~L~~l~~~~~~~~~--~~~~~~~~~~l~~~~~~~~~~f~~v~~~lf~dg   79 (144)
T 2W3L_A          2      DNREIVMKYIHYKLSQRGYEWDAGADSEVVHKTLREAGDDFSRRY--RRDFAEMSSGLHLTPFTARGRFATVVEELFRDG   79 (144)
T ss_dssp               CHHHHHHHHHHHHHHHTTCCCCC---CHHHHHHHHHHHHHHHHHT--HHHHHHHHTTCCCCTTTHHHHHHHHHHHHTTTC
T ss_pred               chHHHHHHHHHHHHHHcCCCCCCCCCChHHHHHHHHHHHHHHHHHh--HHHHHHHHHcCCCChhHHHHHHHHHHHHHhcCC


Q ss_pred                CChHHHHHHHHHHHHHHHHHHhcCChH-HHHHHHHHHHHHHHHHHHHHHHHCCCHHHHHHhhccc
Q Q_BranchBcl2    113    LSWGRVIIAMIFALDVNIKLHGSQTNQ-ENAICSELAELMCAIAGRWLLRRGGWSGMIDAAEPS   175 (197)
Q Consensus       113    inWGRIvalf~F~~~la~~~~~~~~~~~-v~~i~~~~~~fi~~~l~~WI~~~GGW~~~~~~~~~~   175 (197)
                         +|||||+++|+||+.||.+|.+++.+. ++.|++|+++|+.+++.+||.+||||++++++++++
T Consensus       80     inWgrIval~~f~~~la~~~~~~~~~~~v~~i~~~~~~fi~~~l~~WI~~~GGW~~l~~~~~~~   143 (144)
T 2W3L_A          80     VNWGRIVAFFEFGGVMCVESVNREMSPLVDNIALWMTEYLNRHLHTWIQDNGGWDAFVELYGPS   143 (144)
T ss_dssp               CCHHHHHHHHHHHHHHHHHHHHTTCTTHHHHHHHHHHHHCCCCCHHHHHHTTSHHHHHHHTSTT
T ss_pred               CChHHHHHHHHHHHHHHHHHHHcCChHHHHHHHHHHHHHHHHhHHHHHHHCCCHHHHHHHHCCC
```

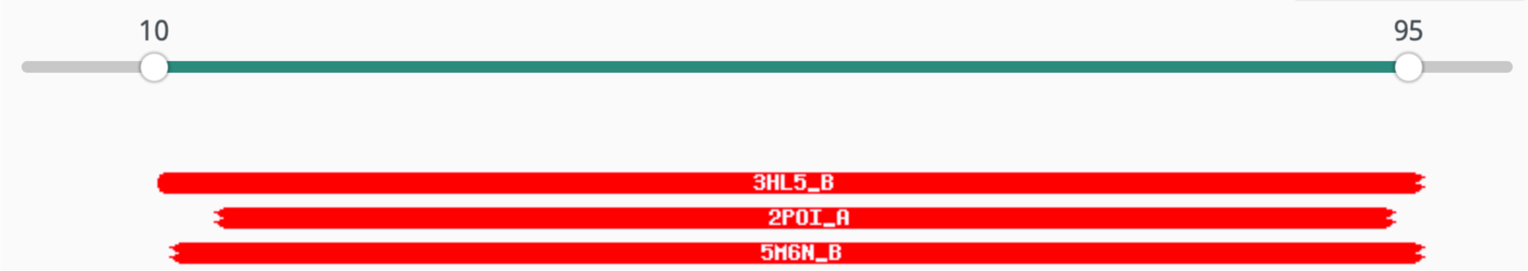


Mytilus IAP

1. **3HL5_B Baculoviral IAP repeat-containing protein 4; BIR, IAP, apoptosis, small molecule; HET: 9JZ; 1.8A {Homo sapiens} SCOP: g.52.1.1; Related PDB entries: 2OPY_A 2JK7_A 5C0K_A 5C0L_A 5C7A_A 5C7B_A 5C7C_A 5C83_A 5C7D_A 5C84_A 5C3H_A 5C3K_A 2OPZ_B 2OPZ_A 2OPZ_D 2OPZ_C 1XB0_A 1XB0_D 1XB0_C 1XB0_E 1XB0_B 1XB0_F 1XB1_A 1XB1_D 1XB1_C 1XB1_E 1XB1_B 1XB1_F 4KMP_B 4KMP_A 3HL5_A 1NW9_A 2VSL_A**
Probability: 99.91%, E-value: 2.5e-27, Score: 148.59, Aligned cols: 84, Identities: 32%, Similarity: 0.578,

```
Q ss_pred              ccccCCCChhhccHHHHHHHhhhCCcccCCCHHHHHHCCEEECCCCCcEEeCccCCeeccCCCCCCHHHHHHHHCCCChh
Q Q_MytilusIAP  10     LFCFNPLHPHMLYYITRQQTYKDWPTQLAQKPADLIRNGFYYTNVGDRVTCFCCGVTLKQWKSSDHIETEHLKYEPNCLF   89 (102)
Q Consensus     10     ~~~~~~~~~~~~~~~~Rl~SF~~wp~~~~~~~~~LA~aGFyy~g~~d~v~Cf~C~~~l~~w~~~d~p~~~H~~~~p~C~f   89 (102)
                       .+...|.++.|..++.|++||.+||..  ++++.||++||||+|..|.|+|++|++.+..|+..++|+.+|++++|+|+|
T Consensus     1      ~~~~~~~~~~~~~~~~Rl~SF~~wp~~--~~~~~LA~aGFyy~g~~d~v~C~~C~~~~~~w~~~d~p~~~H~~~~p~C~f   78 (95)
T 3HL5_B        1      GSHMLPRNPSMADYEARIFTFGTWIYS--VNKEQLARAGFYALGEGDKVKCFHCGGGLTDWKPSEDPWEQHAKWYPGCKY   78 (95)
T ss_dssp             ----CCSCGGGCSHHHHHHHHCSCCSS--SCHHHHHHTTEEECSSTTCEEETTTCCEECSCCTTCCHHHHHHHHCTTCHH
T ss_pred             CCCCCCCChhhcCHHHHHHhhhcCCCC--CCHHHHHHhchhhcCCCCcEEEccCCCeecCCCCCCCHHHHHHHHCCCCHH


Q ss_pred              Hhcccc
Q Q_MytilusIAP  90     AKMVST   95 (102)
Q Consensus     90     ~~~~~~   95 (102)
                       ++..++
T Consensus     79     ~~~~~~   84 (95)
T 3HL5_B        79     LLEQKG   84 (95)
T ss_dssp             HHHHHC
T ss_pred             HHhhcc
```
