## Supplementary material for "Adintoviruses: An Animal-Tropic Family of Midsize Eukaryotic Linear dsDNA (MELD) Viruses": Fig2 Supp3 Additional Adintos

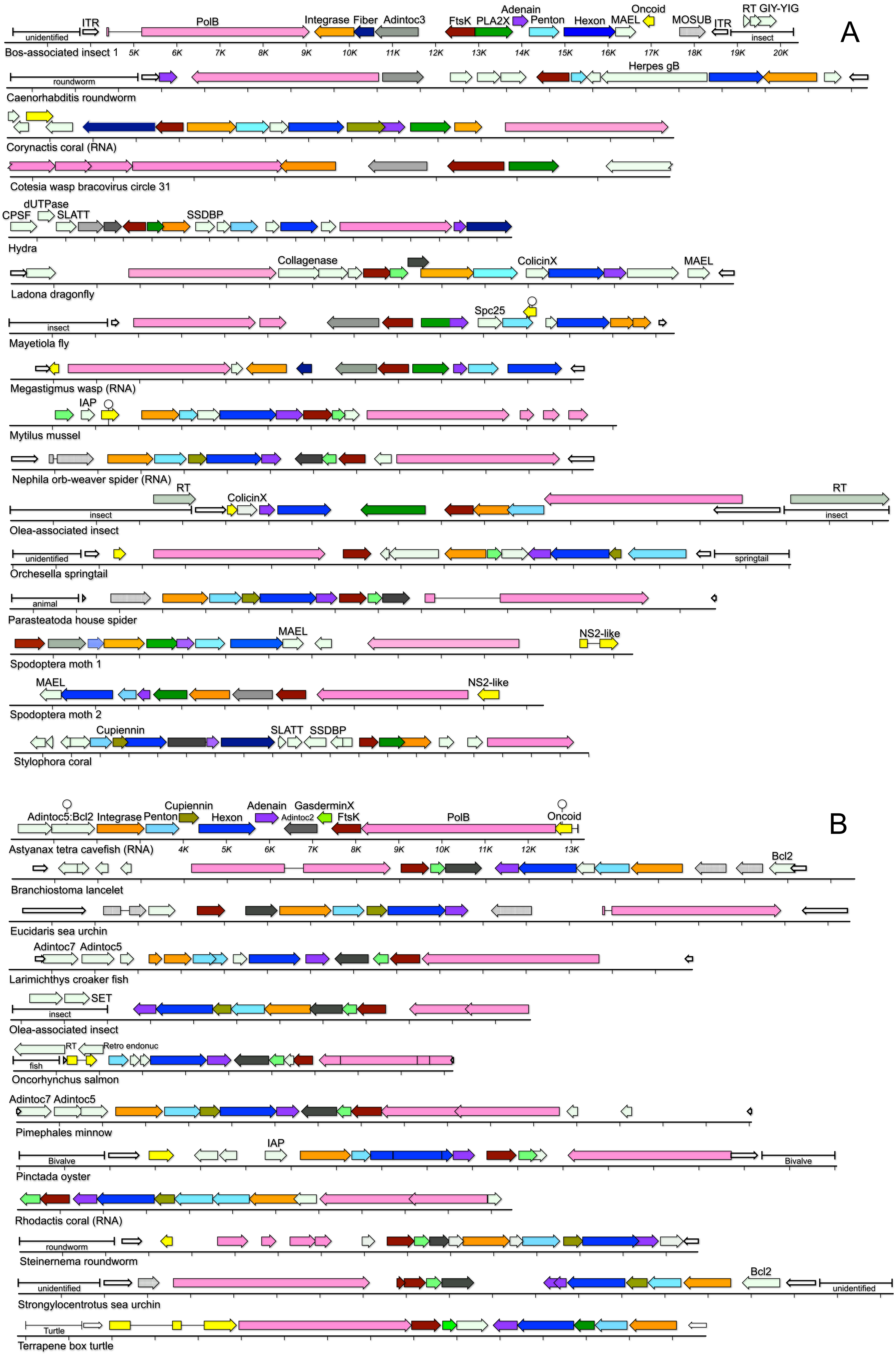


Figure 2 Figure supplement 2: Maps of representative adintovirus contigs recovered from WGS and TSA screens. Panel A shows contigs with Alpha PolB genes. Panel B shows contigs with Beta PolB genes. The figure illustrates that some species, such as the Parasteatoda contig used as bait in the initial screens, unexpectedly unite an Alpha-class PolB with a Beta virion protein operon that includes hallmark Beta-class Cupiennin and GasderminX genes (see Results section "Other adintovirus genes"). Figure 2 Figure supplement 3 lists accession numbers and host taxonomic designations.
